# Horizontal Transfers Lead to the Birth of Momilactone Biosynthetic Gene Clusters in Grass

**DOI:** 10.1101/2022.01.11.475971

**Authors:** Dongya Wu, Yiyu Hu, Shota Akashi, Hideaki Nojiri, Chu-Yu Ye, Qian-Hao Zhu, Kazunori Okada, Longjiang Fan

## Abstract

Momilactone A, an important plant labdane-related diterpenoid, functions as a phytoalexin against pathogens and an allelochemical against neighboring plants. The genes involved in biosynthesis of momilactone A are found in clusters, i.e., MABGCs (Momilactone A biosynthetic gene clusters), in the rice and barnyardgrass genomes. How MABGCs originate and evolve is still not clear. Here, we integrated results from comprehensive phylogeny and comparative genomic analyses of the core genes of MABGC-like clusters and MABGCs in 40 monocot plant genomes, providing convincing evidence for the birth and evolution of MABGCs in grass species. The MABGCs found in the PACMAD clade of the core grass lineage (including Panicoideae and Chloridoideae) originated from a MABGC-like cluster in Triticeae (BOP clade) via horizontal gene transfer (HGT) and followed by recruitment of *MAS* and *CYP76L1* genes. The MABGCs in Oryzoideae originated from PACMAD through another HGT event and lost *CYP76L1* afterwards. The *Oryza* MABGC and another *Oryza* diterpenoid cluster c2BGC are two distinct clusters, with the latter being originated from gene duplication and relocation within Oryzoideae. Further comparison of the expression patterns of the MABGC genes between rice and barnyardgrass in response to pathogen infection and allelopathy provides novel insights into the functional innovation of MABGCs in plants. Our results demonstrate HGT-mediated origination of MABGCs in grass and shed lights into the evolutionary innovation and optimization of plant biosynthetic pathways.

## Introduction

Secondary metabolic compound terpenes play essential roles in plant biotic resistance. At least two biosynthetic gene clusters (BGCs), involved in the biosynthesis of diterpenoid phytoalexins, have been identified in the genome of Asian cultivated rice (*Oryza sativa*). They are c4BGC associated with momilactone A (MA) production (hereafter MABGC) on chromosome 4 and c2BGC for phytocassane production on chromosome 2 (Miyamoto et al., 2016; Toyomasu et al., 2020). Biosynthesis of momilactone in rice requires a series of catalytic reactions, involving enzymes from not only MABGC (CPS4, KSL4, CYP99A2/3, and MAS1/2) but also CYP76M8 from c2BGC, indicating interdependent evolution of the two BGCs (Shimura et al., 2007; Kitaoka et al., 2021) (**Fig. 1a**). A cytochrome P450 enzyme encoded by *CYP701A8* on chromosome 6 is also involved in momilactone biosynthesis in rice (**Fig. 1a**) (Kitaoka et al., 2021). The rice MABGCs were reported to evolve within *Oryza* through duplication and assembly of ancestral biosynthetic genes before the divergence of the BB genome (Miyamoto et al., 2016). MABGCs have also been found in the genomes of paddy weed barnyardgrass *Echinochloa crus-galli* and bryophyte *Calohypnum plumiforme* (Guo et al., 2017; Mao et al., 2020). The functional similarity of MABGCs in grass and bryophyte is likely a result of convergent evolution (Mao et al., 2020; Zhang and Peters, 2020). Compared to the *O. sativa* MABGC, the *E. crus-galli* MABGC has an extra copy of *CYP76L1* (originally wrongly assigned as a member of the CYP76M subfamily) (Guo et al., 2017). Given the divergence time of the two core grass clades, BOP and PACMAD to which rice and barnyardgrass respectively belong, being more than 50 million years ago (Ma et al., 2021) (**Fig. 1b**), hybridization or introgression by sexual or vertical inheritance between the two genera seems illegitimate to result in the patchy occurrence of MABGCs in grass. Hence the origin and evolutionary relationship of MABGCs in grass remains mysterious.

**Figure 1.**
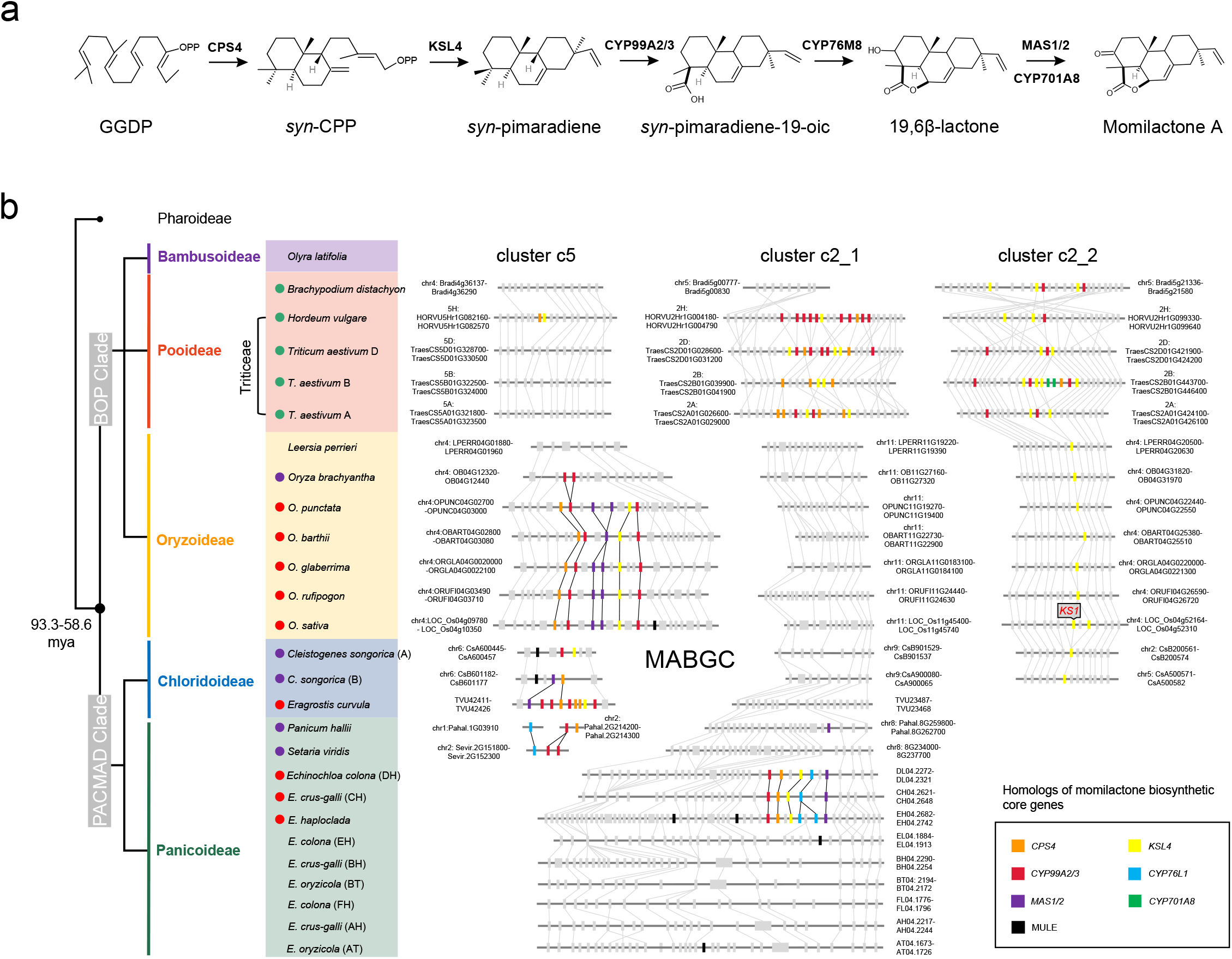
Momilactone biosynthesis pathway and clusters of momilactone A biosynthetic genes (MABGCs) in grass genomes. (**a**) the key steps and corresponding enzymes of the momilactone A biosynthesis process in rice. (**b**) genomic distribution of momilactone biosynthetic genes and their homologs in grass genomes. The left panel shows the phylogenetic topology of the grass family. The red, purple, and green dots in front of the species scientific names represent intact MABGCs, partial MABGCs, and other MABGC-like clusters, respectively. The right panel shows the micro-genomic synteny of MABGC and MABGC-like genes among multi-genomes.

The operon-like gene clustering structure observed in MABGCs is a common phenomenon in bacteria and fungi, in which the origination and evolution of operons have been extensively studied (Nützmann et al., 2018). Gene duplication, neofunctionalization, and relocation are the main routes for cluster formation. Horizontal gene transfer (HGT) occurs widespreadly in prokaryotes and eukaryotes, including grass species (Hibdige et al., 2021). Horizontal transfer is also an important shortcut to the acquisition of gene clusters, particularly in micro-organisms (Slot and Rokas, 2011; Kominek et al., 2019). While high-quality genomes generated in recent years offer good opportunities to trace the evolutionary trajectory of gene clusters in plants (Yang et al., 2021; Liu et al., 2020), we still know little about the mechanisms underlying the birth and evolution of BGCs in plants and whether HGT plays a role in the acquisition of BGCs in grass.

Here, we comprehensively analyzed the individual core genes of MABGCs in 40 monocot genomes using phylogenetics and comparative genomics. We found that it is likely that the grass MABGCs originated from a BGC in Triticeae, which was passed on to the PACMAD and BOP clades via HGT, leading to the formation of the MABGCs observed in rice and barnyardgrass through further gene loss and gain. Our work sheds new insights into evolutionary innovation of momilactone biosynthetic pathway in grass.

## Results and Discussions

### Identification of momilactone biosynthetic genes in grass

Using amino acid sequences of the key MABGC genes (*CPS4, KSL4, CYP99A2/3*, and *MAS1/2*) from rice (*O. sativa*) and *CYP76L1* from barnyardgrass (*E. crus-galli*) as queries, we screened out homologs of all the key MABGC genes from 40 monocot genomes, including 21 species from PACMAD clade (15 in subfamily Panicoideae and 6 in subfamily Chloridoideae), 17 species from BOP clade (2 from Bambusoideae, 8 from Oryzoideae, and 7 from Pooideae), and two outgroup species *Pharus latifolius* and *Ananas comosus* (**Table S1**). The intact MABGCs (defined as harboring at least one copy of *KSL4, MAS1/2, CPS4*, and *CYP99A2/3* or *CYP76L1* homologs within a 200-kb genomic window) were identified in *Oryza* from Oryzoideae, *Echinochloa* from Panicoideae, and *Eragrostis* from Chloridoideae (**Fig. 1b**; **Table S2**). In *Oryza*, while the intact MABGCs were found on chromosome 4 in the AA and BB genomes, only two tandemly duplicated *CYP99A2/3* were found in the FF genome *(Oryza brachyantha),* indicating that MABGCs in *Oryza* had been clustered or generated at least before the divergence of the AA and BB genomes (~6.76 million years ago, mya) (Stein et al., 2018). In *Echinochloa,* MABGCs were found on chromosome 4 in three (sub)genomes (subgenome CH from *E. crus-galli* and DH from *Echinochloa colona* and diploid *Echinochloa haploclada*), which formed a monoclade in *Echinochloa* genome phylogeny (Wu et al., 2021). A candidate MABGC in the genome of weeping lovegrass *Eragrostis curvula* was identified but only partial MABGCs were found in other Chloridoideae species (e.g., *Cleistogenes songorica, Eragrostis nindensis,* and *Eragrostis tef*) (**Fig. 1b**; **Table S2**). Although MABGCs were found in three genera (*Oryza, Eragrostis*, and *Echinochloa*), they are not syntenic in physical genomic positions, implying dynamic evolution of MABGCs in grass (**Fig. 1b**). The dynamic changes of the MABGC loci may be related to the activities of the adjacent mutator-like transposable elements (MuLEs) found near the locations of MABGCs (**Fig. 1b**).

Several MABGC-like clusters were found on chromosomes 2 (clusters c2_1 and c2_2) and 5 (cluster c5) in Pooideae by homology search of MABGC genes (**Fig. 1b**; **Table S2**). These clusters are composed of *CYP99A2/3, KSL4*, and *CPS4* homologs and without *MAS1/2* homologs. Clusters c2_2 are mainly composed of a series of *CYP99A2/3* and *KSL4*, and were assembled before the divergence of Triticeae and *Brachypodium* (**Fig. 1b**). Based on analysis of the syntenic regions in the genomes of Oryzoideae and Chloridoideae, *KSL4* homologs in the c2_2 clusters seem to be derived from tandem duplication of *KS1*, a gene responsible for gibberellin biosynthesis in rice (Toyomasu et al., 2020), and *CYP99A2/3* homologs are embedded among *KSL4* homologs in Pooideae. Intriguingly, the c2_2 cluster in subgenome B of hexaploid *T. aestivum* or tetraploid *T. dicoccoides* contains an additional *CPS4* and two *CYP701A* genes, which are absent in other c2_2 clusters (**Fig.1b**; **Table S2**). Phylogeny and genomic synteny revealed that the two *CYP701A* copies were specifically retained in subgenome B of Triticeae and evolved from the duplication and relocation of the grass-range native *CYP701A* genes (native genes mean a set of conserved syntenic orthologs within highly syntenic blocks from multiple genomes) [e.g., Pl06g08600 from *P. latifolius* (N0), three tandem duplicates Bradi1g37560, Bradi1g37576 and Bradi1g37547 from *B. distachyon* (N1), TraesCS7B01G265800 from *T. aestivum* subgenome B (N2), and CYP701A8 or LOC_Os06g37300 from *O. sativa* (N3) in **Fig. S1**). It should be noted that *CYP701A8* is required for the production of momilactone in rice (*O. sativa*) (Kitaoka et al., 2021) but located away from the MABGC, while in the cluster c2_2 of *T. aestivum* subgenome B, two *CYP701A8* homologs were found in the MABGC-like cluster. Cluster c5 composed of *CPS4* and *KSL4* homologs was found only in the genome of *H. vulgare* (**Fig.1b**). Cluster c2_1 is absent in the *Brachypodium* genome, implying a recent origination of c2_1 within Triticeae. In *T. aestivum*, while the cluster c2_1 of subgenome B has no *CYP99A2/3*, both c2_1 clusters of subgenomes A and D contain at least one copy of *KSL4, CPS4*, and *CYP99A2/3* (**Fig. 1b**).

To know whether these MABGC-like clusters function in plant pathogen resistance, expression profiling of the genes of those clusters was investigated for their responses to infection of different pathogens in *T. aestivum* (**Table S3**). The expression level of two *KSL4*, one *CYP99A2/3*, two *CYP701A8,* and one *CPS4* from the cluster c2_2 of subgenome B increased dramatically in response to infection of the pathogen of *Fusarium* head blight, crown rot, powdery mildew, or tan spot, while almost no expression change was observed for the genes of the same cluster from subgenomes A and D. Comparing to cluster c2_2, the three c2_1 clusters from the three subgenomes showed opposite response upon infection of these pathogens, where the clusters of subgenomes A and D exhibited significant expression increase, while no response was observed for c2_1 of subgenome B (**Table S3**). Comparing the cluster components of the three *T. aestivum* subgenomes, c2_1 of subgenome B harbors additional *CPS4* and *CYP701A* genes and c2_2 of subgenome B includes no *CYP99A2/3* (**Fig. 1b**). Opposite responses of the clusters c2_2 and c2_1 from subgenome B and of those from subgenomes A and D upon infection of the same pathogens suggest functional redundancy of the two clusters, leading to differential loss of core genes of the clusters in different subgenome and to retaining only a single copy of intact functional cluster (with *CPS4, KSL4*, and *CYP99A2/3*). In addition, within each functional cluster (c2_2 of subgenome B and c2_1 of subgenomes A and D), only a sub-cluster of genes functioned in response to pathogen infection (**Table S3**). For example, of the 10 genes of the cluster c2_2 of subgenome B, only six continuously distributed genes (two *KSL4*, two *CYP701A*, one *CPS4*, and one *CYP99A2/3* homologs) responded to pathogen infection. Taken together, the MABGC-like clusters in wheat showed subgenome-biased response to a range of pathogen stresses.

### Horizontal transfer of MABGC genes among grass

The divergence time between the BOP and PACMAD clades is more than 50 mya (Ma et al., 2021). The disperse distribution of MABGCs and MABGC-like clusters in different grass species evokes the wonders of the evolutionary relationship among these clusters. Previous study suggested that MABGCs in *Echinochloa* and *Oryza* arose independently by acquisition of core enzymes (Mao et al., 2020; Zhang and Peters, 2020). Whether the MABGC-like clusters in Triticeae are related to the origin of MABGCs has never been investigated. Thus, we performed comprehensive phylogenetic analyses across the whole grass family to infer the evolution of the core biosynthetic genes in these clusters and to trace the trajectory of MABGC origination.

### Evolution of CPS4

CPS4, *syn*-copalyl diphosphate synthase 4, catalyzes GGDP into *syn*-CPP, the first reaction of the momilactone biosynthesis pathway (**Fig. 1a**). Based on the phylogeny tree, the *CPS4* homologs in MABGCs of *Oryza, Echinochloa,* and *Eragrostis* were clustered in a monoclade (MABGC clade), nested within *CPS4* homologs from Pooideae (**Fig. 2a**). The *CPS4* homologs in Pooideae were mainly assigned separately in three lineages, corresponding to clusters c2_1, c2_2, and c5, of which the *CPS4* homologs from cluster c2_2 is located at the basal position. *CPS2* and *CPS3* were identified as homologs of *CPS4* in rice. Except the homologs from the MABGC clade and *CPS3* clade, the *CPS4* homologs in grass were found to have a congruent relationship as the species phylogeny, thus homologs at the subfamily level were grouped together and have conserved physical positions among Poaceae genomes (**Fig. 2b**). *CPS2* from *O. sativa* (N0 in **Fig. 2b**, LOC_Os02g36210) is syntenic to Pl02g21350 (N1) from *P. latifolius* (Pharoideae, basal group in grass family), Zlat_10013826 (N3) from *Zizania latifolia* (Oryzoideae), LPERR02G17320 (N4) from *Leersia perrieri* (Oryzoideae), Ola021376 (N2) from *Olyra latifolia* (Bambusoideae), Et_1A_005482 (N5) from *Eragrostis tef* subgenome A (Chloridoideae), and eh_chr7.2132 (N6) from *Echinochloa haploclada* (Panicoideae). Therefore, both phylogeny and genomic synteny indicate that the *CPS4* homologs of the MABGC monoclade are neither native copies nor duplicated paralogs of native copies (e.g., *CPS2* in *O. sativa*) but instead were highly possibly originated by HGT. It was noticed that *CPS3* genes in *Oryza* were nested in PACMAD lineage, implying the possibility of another HGT event. To test the HGT hypothesis, topology tests (including bp-RELL, KH, SH, and ELW tests) on constrained trees were employed to determine whether the HGT tree (tree 1 in **Fig. 2c**) could statistically explain the data better than non-HGT (native origin, tree 2 and tree 3) phylogenies. Results showed that the HGT phylogeny of *CPS4* was strongly supported and non-HGT phylogenies were rejected (test 1 in **Fig. 2c**). Within the MABGC clade, *CPS4* genes from three subfamilies (Oryzoideae, Panicoideae, and Chloridoideae) form three distinct groups (**Fig. 2a**). We constructed constrained trees to test the topology robustness using Pooideae *CPS4* homologs as outgroup, and neither of three topologies could be rejected (test 2 in **Fig. 2c**).

**Figure 2.**
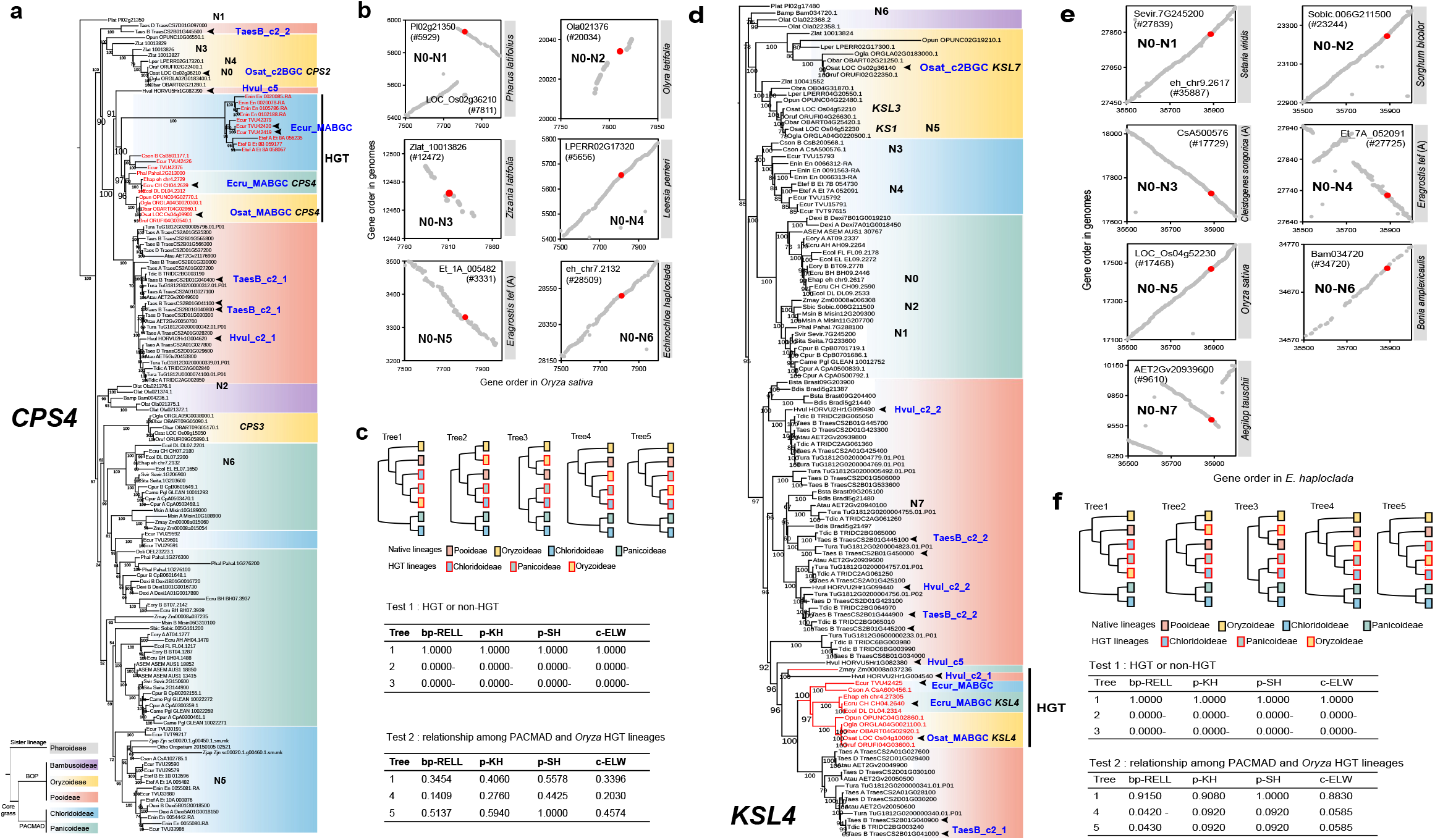
Phylogeny and genomic synteny analyses of *CPS4* and *KSL4* genes. (**a**) the maximum-likelihood tree of *CPS4* and its homologs across the grass family. The homolog in *P. latifolius* was set as an outgroup. Different background colors represent different subfamilies. (**b**) genomic synteny among the conserved native *CPS4* homologs. Red dots represent that the two *CPS4* homologs from two genomes are in good synteny. (**c**) topology tests on five constrained trees. The top panel shows the topologies of constrained trees used in tests. The middle (Test 1) and bottom (Test 2) panels show the results of the two tests on the HGT event of *CPS4* from Pooideae to PACMAD and the HGT from PACMAD to *Oryza*, respectively. (**d**) the phylogenetic tree of *KSL4* and its homologs. (**e**) genomic synteny among the conserved native *KSL4* homologs. Red dots represent that the two *CPS4* homologs from two genomes are in good synteny. (**f**) topology tests on five constrained trees to test the robustness of the hypothesis of HGT for *KSL4*.

### Evolution of KSL4

KSL4, *ent*-kaurene synthase 4, cyclicizes *syn*-CPP into *syn*-pimaradiene (**Fig. 1a**). *KSL4* genes from MABGCs form a highly supported monoclade nested within the Triticeae lineage (**Fig. 2d**). The clade composed of *KSL4* homologs from the c2_2 clusters of Pooideae is located at the base within the Pooideae lineage, and the *KSL4* clade from the c2_1 clusters is a sister to the MABGC clade. The genomic positions of tandem duplicates *KS1* and *KSL3* from rice c2BGC are highly conserved in grass family with perfect genomic synteny among grass genomes (**Fig. 2e**). *KS1* (N5, LOC_Os04g52230) or *KSL3* (LOC_Os04g52210) from *O. sativa* (Oryzoideae), Bam034720 from *Bonia amplexicaulis* (Bambusoideae), AET2Gv20939600 from *Aegilop tauschii* (Pooideae), CsA500576 from *Cleistogenes songorica* subgenome A (Chloridoideae), Et_7A_052091 from *E. tef* subgenome A (Chloridoideae), Sobic.006G211500 from *Sorghum bicolor* (Panicoideae), and Sevir.7G245200 from *Setaria viridis* (Panicoideae) are all syntenic to the native *KSL4* homolog eh_chr9.2617 (N0) from *E. haploclada* (Panicoideae). *KSL7* was integrated into rice c2BGC as a duplicate of *KS1/KSL3*. These results indicate that *KS1* or *KSL3* is a native conserved *KSL* gene in grass family, with the fundamental function of *KS1* in synthesizing gibberellins (Miyamoto et al., 2016), and *KSL4* is probably inherited from Triticeae by HGT, rather than gene duplication, convergent evolution or in-complete lineage sorting. Topology tests strongly support the HGT origination of *KSL4* (test 1 in **Fig. 2f**). Within the HGT lineage, three clades from three subfamilies are formed. Topology tests based on bp-RELL rejected the topology that *KSL4* lineage in *Oryza* is sister to the common ancestor of Panicoideae and Chloridoideae lineages (test 2 in **Fig. 2f**), indicating that *KSL4* in *Oryza* was derived from Panicoideae or Chloridoideae.

### Evolution of CYP99A2/3

CYP99A2/3 functions as a C19 oxidase (**Fig. 1a**). The *CYP99A2/3* homologs are separated into two main lineages in the phylogeny tree, consistent with species phylogeny (PACMAD and BOP clades) (**Fig. S2a**). In the BOP lineage, two monoclades (HGT1 and HGT2) are nested within *CYP99A2/3* homologs from Pooideae. Monoclade HGT2 composed of genes from Panicoideae is a sister to one clade from Pooideae. Monoclade HGT1, including *CYP99A2/3* genes of MABGCs, composed of three sub-clades from *Oryza*, Panicoideae and Chloridoideae, is a sister to the clade containing Triticeae clusters c2_1 (**Fig. S2a**). *CYP99A2* and *CYP99A3* in *Oryza* arose from tandem duplication after the divergence of the AA/BB and FF *Oryza* genomes. Besides *Echinochloa*, *CYP99A2/3* genes were found in other species in Panicoideae and especially expanded in *Setaria.* In Chloridoideae, the copies of *CYP99A2/3* were expanded in *Eragrostis*. Topology tests strongly support that clade HGT1 arose from Pooideae lineage (test 1 in **Fig. S2b**), but the phylogenetic relationship among clades from three subfamilies within the HGT1 clade could not be deciphered distinctly (test2 in **Fig. S2b**).

### Evolution of MAS1/2

MAS1/2 catalyzes the oxidation of 3β-hydroxy-syn-pimaradien-19,6β-olide to form the characteristic C3 keto group in rice (**Fig. 1a**). *MAS1* and *MAS2* genes are tandem duplicates in *Oryza*, nested within PACMAD linage, and are sisters to the clade from Chloridoideae (**Fig. S3a**). *MAS3* genes in grass form a congruent topology just like the subfamily-level species phylogeny. Their genomic positions are conserved across grass genomes (**Fig. S3b**). The conserved homologs of *MAS1/2* are absent in Pooideae and Bambusoideae, indicating that *MAS1/2* homologs are specific for the PACMAD clade. Topology tests revealed that *MAS1/2* in *Oryza* were derived from PACMAD via HGT (**Fig. S3c**).

Based on above phylogenetic and comparative genomics analyses of the core genes of MABGCs across grass genomes, we propose that the genomic position of *KSL4* homolog in the c2_2 cluster of Pooideae, which is syntenic to *KS1* in rice, is the eventual origin of MABGC-like clusters in Pooideae, such as cluster c5 and cluster c2_1 formed by assembly of *KSL4* homologs with *CPS4* and *CYP99A2/3* (**Fig. 1b**). MABGC-like cluster was then passed on to the common ancestor of Panicoideae and Chloridoideae by HGT, where the MABGC with indispensable core genes required for momilactone biosynthesis was formed by further integration of *MAS1/2* (**Fig. 3**). The *Oryza* MABGCs were acquired from Panicoideae or Chloridoideae via another HGT event because the genes from the *Oryza* MABGC are sisters to or nested within PACMAD homologs and topology tests (bp-RELL) on *KSL4* and *MAS1/2* homologs rejected the topology that *Oryza* genes are sisters to common ancestors of Panicoideae and Chloridoideae (**Figs. 2f and 3**; **Fig. S3c**). HGT is commonly observed in grass species (Hibdige et al., 2021). Occasional physical contacts (e.g., illegitimate hybridization or grafting) or intermediation by parasitic plants (e.g., *Striga* and *Aeginetia*), insects or pathogens are potential explanations for genetic material transfer between BOP and PACMAD clade (Xia et al., 2021; Wang et al., 2020).

**Figure 3.**
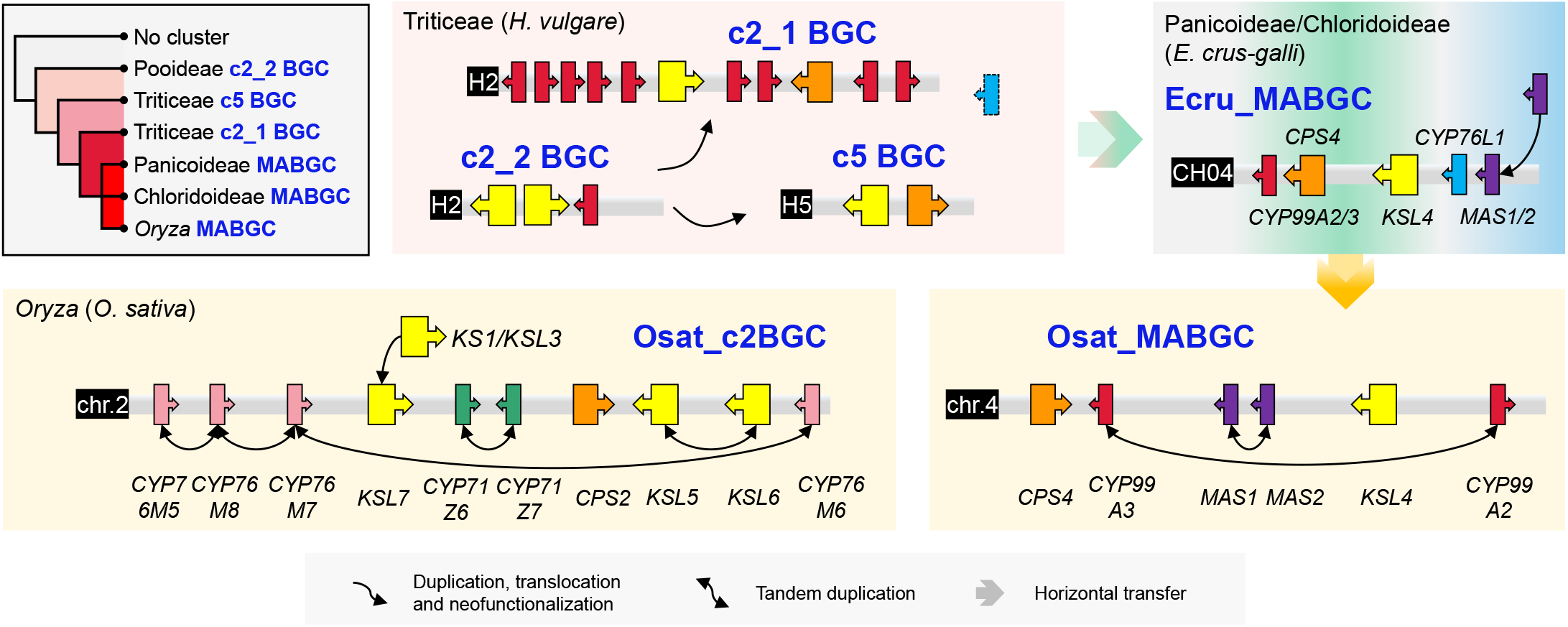
The evolutionary trajectory of MABGCs in grass. The biosynthetic gene cluster (BGC) structures were shown using representative species (*H. vulgare* from Pooideae or Triticeae, *E. crus-galli* from PACMAD clade, *O. sativa* from *Oryza* or Oryzoideae). Cluster c2_2 in Pooideae or Triticeae is the eventual origin of MABGCs. Clusters c2_1 and c5 arose through gene duplication and translocation of cluster c2_2. Subsequently the ancient c2_1 or c5 (MABGC-like) was transferred into the PACMAD clade via HGT, where the MABGC-like recruited *CYP76L1* and *MAS1/2,* leading to the birth of MABGC. *Oryza* species acquired the PACMAD clade MABGC via HGT, followed by loss of *CYP76L1.* The *Oryza* MABGC is not the duplicate of Osat_c2BGC, another diterpenoid BGC. Tandem duplication happened during the evolution of MABGC and c2BGC in rice.

Different to the origination of MABGC, c2BGC of Oryzoideae was formed via gene duplication, translocation and neofunctionalization rather than HGT. *CPS2* is conserved across the whole grass family (**Figs. 2a** and **2b**) and is the ancestor of *CPS4* of MABGC. *KSL7* was translocated along with *CPS2* as a duplicate of *KS1/KSL3* (**Fig. 3**). *CYP76M7* and *CYP76M8* were recently duplicated in *Oryza* (**Fig. S4**), although *CYP76M7* plays roles in phytocassane biosynthesis and *CYP76M8* is mainly responsible for momilactone production (Kitaoka et al., 2021). *CYP71Z6* and *CYP71Z7* are tandem duplicates, phylogenetically neighboring to four *CYP76Z* genes from c7BGC (a cluster on chromosome 7, associated with production of the casbane-type diterpenoid phytoalexin ent-10-oxodepressin) (Liang et al., 2021) (**Fig. S5**). *KSL5* and *KSL6* are *Oryza*-specific tandem duplicates, neighboring to *KSL8* (responsible for the biosynthesis of Oryzalexin S) and *KSL10* (responsible for the biosynthesis of Oryzalexins A-F) (Nemoto et al., 2004; Miyamoto et al., 2016) (**Fig. S6**).

### Comparison of MABGCs between *Oryza* and *Echinochloa* species

Comparing to *Oryza* MABGCs, *Echinochloa* MABGCs have an extra copy of cytochrome P450 gene *CYP76L1* (**Fig. 1b**). The phylogenetic tree of *CYP76L1* homologs is composed of two major lineages, in line with species phylogeny (BOP and PACMAD lineage) (**Fig. S7a**). *CYP76L1* genes of MABGCs are nested within the Pooideae lineage and also found in *Setaria* and *Panicum.* Genome synteny was used to rule out the possibility of convergent evolution and incomplete lineage sorting (ILS) to cause the discordance between gene topology and species phylogeny. Taking *O. sativa* genome as a reference, we scanned the genomic synteny around the native *CYP76L1* regions among grass genomes (**Fig. S7b**). Lper_LPERR09G08730 (N1) in *Leersia perrieri* (Oryzoideae), eh_chr3.2603 (N2) in *E. haploclada* (Panicoideae), Et_2A_016179 (N3) in *E. tef* subgenome A (Chloridoideae), Ola025389 (N4) from *O. latifolia* (Bambusoideae), and AET5Gv20559300 (N5) from *A. tauschii* (Pooideae) are all highly syntenic to LOC_Os09g27500 (N0) from *O. sativa*, indicating that Paniceae *CYP76L1* genes nested within Pooideae genes in the gene tree were acquired via HGT as extra copies of the native *CYP76L1.* Topology tests strongly support the HGT origination of *CYP76L1* in MABGCs from Pooideae (**Fig. S7c**). To distinguish the two types of *CYP76L1,* we named the native *CYP76L1* homologs and *CYP76L1* acquired by HGT as CYP76L1-native and CYP76L1-hgt, respectively. CYP76L1-native (LOC_Os9g27500), but not CYP76L-hgt, is present in the *O. sativa* genome, while both CYP76L1-native and -hgt are present in barnyardgrass MABGC (**Fig. 1b**; **Fig. S7a**). Given the presence of *CYP76L1* in barnyardgrass MABGC and its strong HGT signal based on phylogenetic tree, we speculate that *CYP76L1* was transferred from Pooideae to PACMAD along with MABGC core genes as a cluster, and subsequently lost in *Oryza* (**Fig. 3**).

In barnyardgrass (*E. crus-galli*), CYP76L1-hgt is co-expressed with other MABGC genes (Sultana et al., 2019). A recent study in rice demonstrated that CYP76L1-hgt from barnyardgrass is functionally equivalent to *CYP76M8* in catalyzing the 6β-hydroxylation of syn-pimaradiene (Kitaoka et al., 2021), implying a function of CYP76L1-hgt in momilactone biosynthesis. CYP76L1-native may not play a role in momilactone production. First, gene co-expression network analysis revealed that CYP76L1-native is not a component of the momilactone production pathway in rice (**Fig. 4a**). Second, upon treatment of jasmonic acid, the content of momilactone in rice roots was highly induced (Kitaoka et al., 2021), but no up-regulation of CYP76L1-native was found (**Fig. S8a**).

**Figure 4.**
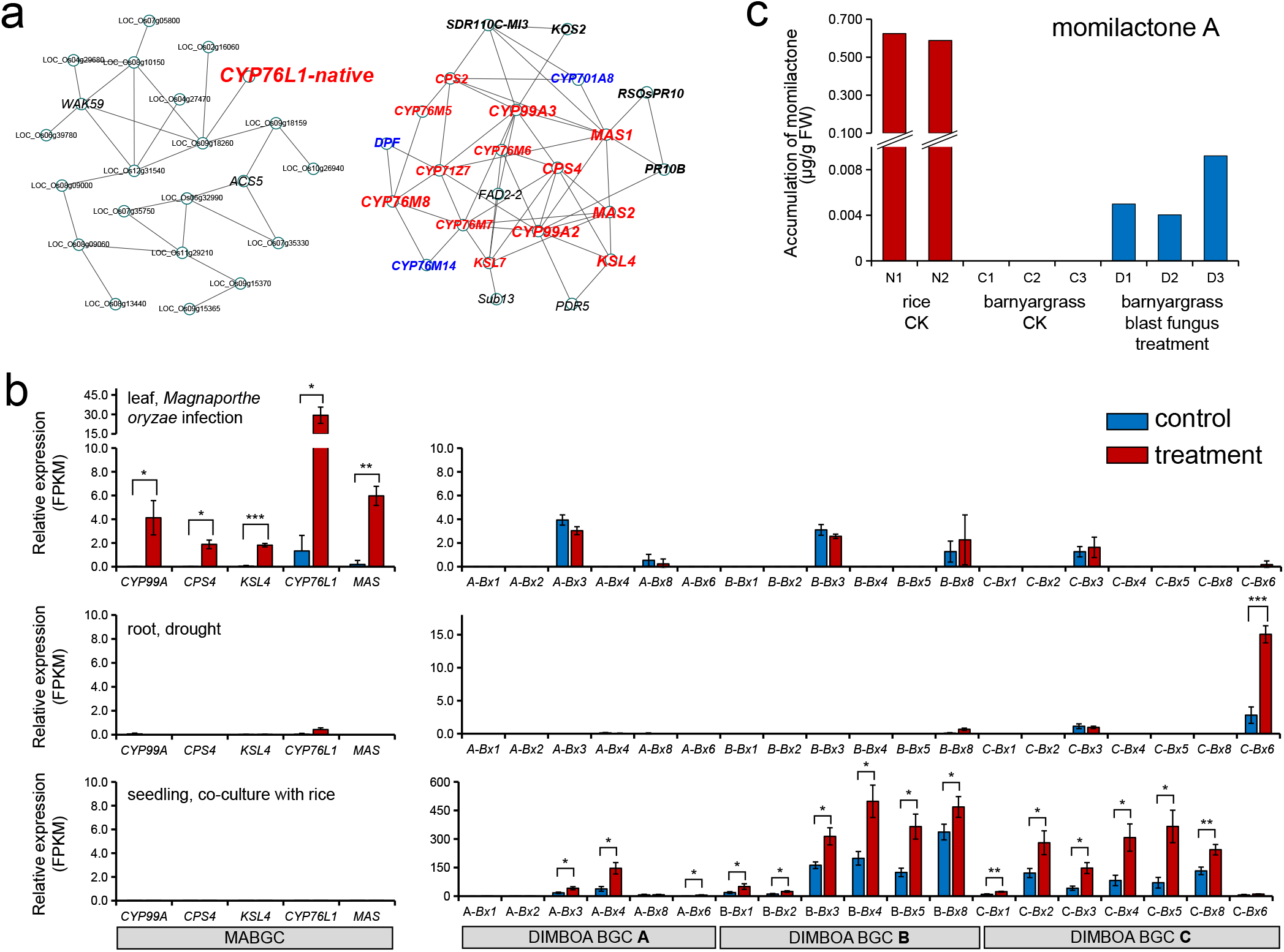
Comparison of MABGCs from rice (*O. sativa*) and barnyardgrass (*E. crus-galli*). (**a**) gene co-expression network of native *CYP76L1* (CYP76L1-native) reveals the independent relationship between *CYP76L1*-native and the momilatone biosynthetic genes in rice. (**b**) transcriptomic profiles of barnyardgrass MABGC and the gene cluster for DIMBOA (Bx cluster) in response to *M. oryzae* infection, drought, or co-culture with rice. *, *P*<0.05; **, *P*<0.01; ***, *P*<0.001, Student’s *t* test. (**c**) Comparison of the content of momilactone A in leaves of rice and barnyardgrass between control (CK) and *M. oryzae* infection. N1 to N2, C1 to C3, and D1 to D3 are biological replicates of the three experiments.

Infection by blast pathogen *M. oryzae* induces the production of momilactone in rice (Hasegawa et al., 2010), likely due to up-regulation of MABGC genes (except *MAS2*) (**Fig. S8b**). In barnyardgrass (*E. crus-galli*), consistent with significant up-regulation (*P* < 0.05) of MABGC genes, including CYP76L1-hgt (**Fig. 4b**), the content of momilactone A was also significantly induced by *M. oryzae* infection, but its level was ~100-times lower than that observed in rice plants without pathogen infection (**Fig. 4c**). Copy number variations in *MAS1/2* and *CYP99A2/3* between the two species may contribute to biosynthesis and accumulation of momilactone, and the catalytic efficiency difference between *CYP76M8* and CYP76L1-hgt may be another contributor for the difference.

Remarkably, barnyardgrass is one of the most detrimental weeds in paddy fields. Momilactone secreted by rice roots plays a critical role in rice allelopathy, by which growth of barnyardgrass in the neighboring environments of rice plants but not the rice plants themselves is inhibited by momilactone because barnyardgrass is much more sensitive to momilactone than rice (Kato-Noguchi and Peters, 2013). Barnyardgrass induces rice to secrete momilactone but the barnyardgrass-induced rice allelopathy is not associated with the MABGC of barnyardgrass as no gene from the barnyardgrass MABGC was induced under the rice and barnyardgrass co-planting conditions (**Fig. 4b**). Instead, genes of the barnyardgrass DIMBOA cluster, involved in the production of benzoxazinoids secondary metabolites, were significantly up-regulated under the co-planting conditions (**Fig. 4b**) (Guo et al., 2017). While the DIMBOA cluster is only ~359 kb away from the MABGC in the *E. crus-galli* genome (**Fig. S9**), the two barnyardgrass BGCs are not co-expressed (Sultana et al., 2019) and have distinct functions based on their expression profiles responding to pathogen infection and co-cultivation with rice (**Fig. 4b**), with the MABGC being involved in response to pathogen infection and the DIMBOA cluster being involved allelopathic interaction.

## Conclusion

By integrating gene phylogeny and comparative genomics analyses using 40 monocot genomes, we show that intact MABGCs are present in *Oryza* from Oryzoideae (BOP clade), *Echinochloa* from Panicoideae (PACMAD clade), and *Eragrostis* from Chloridoideae (PACMAD clade), and revealed the evolution trajectory of MABGCs in grass, i.e., MABGCs in PACMAD had arisen from horizontal gene transfer from MABGC-like clusters in Triticeae followed by further integration of *MAS* genes, which was then acquired by Oryzoideae via another HGT event. Our study also demonstrate functional innovation of MABGCs in plants.

## Materials and Methods

### Homolog Identification of Biosynthetic Genes

The protein sequences of momilatone and phytocassane biosynthetic genes of *O. sativa* (*CPS4*, LOC_Os04g09900; *KSL4*, LOC_Os04g10060; *CYP99A2/3*, LOC_Os04g09920 and LOC_Os04g10160; *MAS1/2*, LOC_Os04g10000 and LOC_Os04g10010; *KSL5/6,* LOC_Os02g36220 and LOC_Os02g36264; *CYP76M5/6/7/8*, LOC_Os02g36030, LOC_Os02g36280, LOC_Os02g36110 and LOC_Os02g36070; *CYP71Z6/7*, LOC_Os02g36150 and LOC_Os02g36190; *CYP701A8*, LOC_Os06g37300) and of *E. crus-galli* (*CYP76L1*, CH04.2641) were set as reference baits. For the genes with multiple transcripts, the longest ones were selected. BLASTP was performed using the reference baits against more than 1.95 million protein sequences from the genomes of 39 grass species and *A. comosus* (**Table S1**). Raw homologs were selected with the filtering criteria of BLAST *e*-value less than 1e-30 and identity greater than 50%. Raw homologs were aligned using MAFFT (v7.310) (Katoh and Standley, 2013) and the phylogenetic trees were built using IQ-TREE (v1.6.6) under the best substitution model with 1000 replicates for bootstrap (Nguyen et al., 2015). Using *A. comosus* and *P. latifolius* as outgroup species, the final repository of MABGC homologs were determined according to the phylogeny and used in the following analysis.

### Phylogeny Analysis

Homologous sequences of each biosynthetic gene were re-aligned using MAFFT (v7.310) and trimmed using Gblocks (v0.91b) with the parameter “-b4=5 -b5=h” (Castresana, 2000). The best substitution model for each trimmed alignment was determined by ModelFinder and the phylogenetic tree was constructed by IQ-TREE (v1.6.6) for 1000 bootstrap replicates (Nguyen et al., 2015). Constrained trees were searched and built in IQ-TREE (v1.6.6) and topology tests on them were performed with 10000 times of bootstrapping, including bootstrap proportion (BP) test, Kishino-Hasegawa (KH) test, Shimodaira-Hasegawa (SH) test, and expected likelihood weights (ELW).

### Genomic Synteny

Protein sequences from the grass genomes were compared pairwise using BLASTP and the best hits with thresholds of *e*-value less than 1e-30 and identity greater than 50% were kept. According to their physical positions in each genome, the genes were ordered. Each blast best-hit had a pair of coordinates and the gene-to-gene synteny was plotted pairwise for grass genomes.

### Gene Expression Analysis

For hexaploid wheat (*T. aestivum*) gene expression profiling, the quantified expression values of genes in different tissues and under different pathogen infections (*Fusarium* head blight pathogen *Fusarium graminearum*, crown rot pathogen *Fusarium pseudograminearum*, powdery mildew pathogen *Blumeria graminis*, tan spot pathogen *Pyrenophora tritici-repentis*, stripe rust pathogen *Puccinia striiformis*, and black chaff pathogen *Xanthomonas translucens*) were obtained from WheatOmics 1.0 database (http://wheatomics.sdau.edu.cn/)(Ma et al., 2021). For barnyardgrass (*E. crus-galli*), the RNA-seq datasets under the treatment of co-culture with rice seedling (Guo et al., 2017), infection by blast fungus *M. oryzae* and drought induced by polyethylene glycol (PEG) (Ye et al., 2020) were mapped against the reference genome STB08 (Wu et al., 2022) and quantified using TopHat (v2.1.1) and Cufflinks (v2.2.1) (Trapnell et al., 2012). For rice (*O. sativa*), the co-expression network was built in RiceFREND database (https://ricefrend.dna.affrc.go.jp/) (Sato et al., 2013a) and the expression data from samples subjected to jasmonic acid (JA) treatment, drought stress, and blast fungus *M. oryzae* infection were obtained from RiceXPro (https://ricexpro.dna.affrc.go.jp/) (Sato et al., 2013b) and Plant Public RNA-seq Database (http://ipf.sustech.edu.cn/pub/ricerna/).

### Momilactone Quantification

*Echinochloa* leaves infected by blast fungus or mock control were prepared before the analysis. Each treated sample (roughly 100 mg) was submerged in 4 ml of 80% methanol at 4°C for 24 h, and 5 μL of the extract was subjected to LC-MS/MS analysis using API-3000 with an electrospray ion source (Applied Biosystems Instruments, Foster City, CA, USA) and an Agilent 1100 HPLC instrument (Agilent Technologies, Palo Alto, CA, USA) equipped with a PEGASIL C18 column (150 mm long, 2.1 mm in diameter; Senshu Scientific, Tokyo, Japan) with the selected reaction monitoring transitions (for momilactone A, m/z 315→ 271) as described previously (Miyamoto et al., 2016).

## Acknowledgement

This work was supported by the National Natural Science Foundation (31971865 and 32170621), the Department of Science and Technology of Zhejiang Province (2022C02032), Zhejiang Natural Science Foundation (LZ17C130001) and Jiangsu Collaborative Innovation Center for Modern Crop Production. The authors declare no competing interests.

## Author contributions

L.F. and D.W. conceived the research. D.W. and Y.H. performed the data analysis. S.A., H.N. and K. O. performed the experimental quantification of momilactones in rice and barnyardgrass. C.-Y.Y., K.O., Q.-H.Z. and L.F. discussed the findings. Q.-H.Z. edited the manuscript. D.W. and L. F. wrote the manuscript. All authors read and contributed to the manuscript.

## Supplementary Information

**Table S1.** A list of plant genomes used in this study.

**Table S2.** Homologs of the key genes (*CPS4/KSL4/CYP99A/CYP76L1/MAS/CYP701A8*) involved in momilactone biosynthesis in grass species.

**Table S3.** Transcriptomic profiling of wheat (*T. aestivum*) under infection of pathogens.

**Figure S1.** Phylogeny and genomic synteny of *CYP701A8* homologs in grass. Genes in different subfamilies are marked in different colored backgrounds. Genes used in synteny analyzes are marked in the phylogenetic tree from N0 to N3. Homologs in Triticeae subgenomes B and four CYP701A genes in *O. sativa* are highlighted. The red syntenic dots represent pairs of native homologs between genomes.

**Figure S2.** Phylogeny of *CYP99A2/3* homologs in grass and topology tests. (**a**) Homolog phylogeny of *CYP99A2/3*. Genes in different subfamilies are marked in different colored backgrounds. Cluster information of some homologs in Triticeae and MABGCs are suggested. (**b**) Topology tests. Three and three constrained trees were set for test1 and test2, respectively. Minus signs represent that the corresponding topology could be rejected significantly.

**Figure S3.** Phylogeny and genomic synteny of *MAS1/2* homologs in grass and topology tests. (**a**) The maximum-likelihood tree of *MAS1/2* and its homologs across the grass family. The homolog in *A. comosus* was set as an outgroup. Different background colors represent different subfamilies. (**b**) Genomic synteny among the native *MAS3* homologs. Red dots represent that the two *MAS3* homologs from two genomes are in good synteny. (**c**) Topology tests on three constrained trees. The top panel shows the topologies of constrained trees used in tests. The bottom panels show the results of the test on the HGT event of *MAS1/2* from PACMAD to *Oryza*. Minus signs represent that the corresponding topology could be rejected significantly.

**Figure S4.** A maximum-likelihood phylogenetic tree of *CYP76M5/6/7/8* homologs in grass. Different background colors represent different subfamilies. The branch containing *CYP76M5/6/7/8* from O. sativa is zoomed in.

**Figure S5.** A maximum-likelihood phylogeny of *CYP71Z6/7* homologs in grass. Different background colors represent different subfamilies. The branch containing *CYP71Z* genes from *O. sativa* is zoomed in.

**Figure S6.** A maximum-likelihood phylogeny of *KSL5/6* homologs in grass. Different background colors represent different subfamilies.

**Figure S7.** Phylogeny and genomic synteny of *CYP76L1* homologs in grass and topology tests. (**a**) The maximum-likelihood tree of *CYP76L1* and its homologs across the grass family. Different background colors represent different subfamilies. Genes used in synteny analyzes are marked in the phylogenetic tree from N0 to N5. (**b**) Genomic synteny among the native *CYP76L1* homologs. Red dots represent that the two homologs from two genomes are in good synteny. (**c**) Topology tests on constrained trees. The top panel shows the topologies of constrained trees used in tests. The bottom panels show the results of test on the HGT of *CYP76L1* from Pooideae to Panicoideae. Minus signs represent that the corresponding topology could be rejected significantly.

**Figure S8.** Expression of MABGC, c2BGC and related genes in rice under JA treatment (**a**), drought and *M. oryzae* infection (**b**).

**Figure S9.** The physical positions of MABGCs and DIMBOA Bx clusters in *Echinochloa* genomes.

