## Supplementary material for "Horizontal Transfers Lead to the Birth of Momilactone Biosynthetic Gene Clusters in Grass"

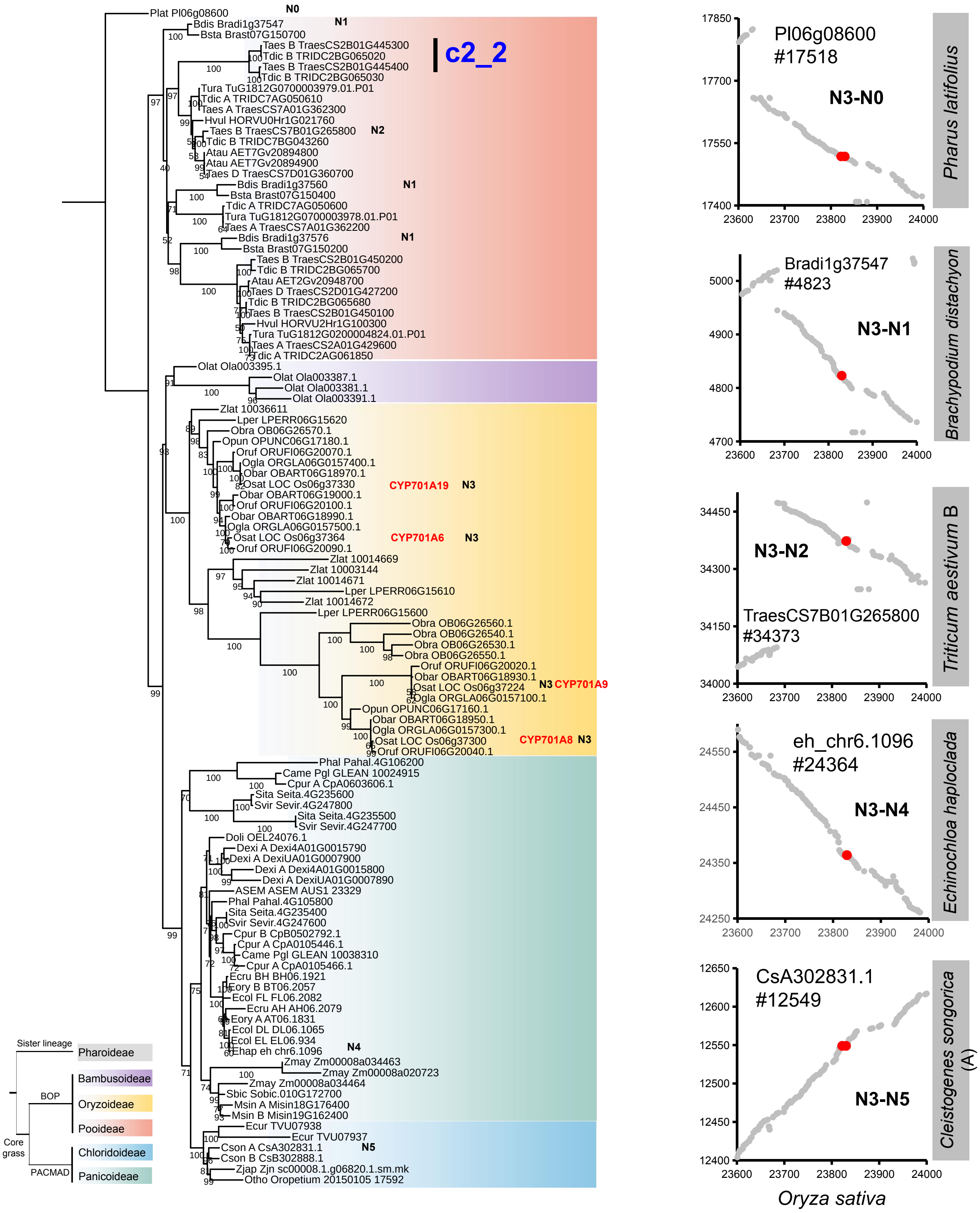

c2_2
N0
N1
N2
N3
N4
N5
CYP701A19
CYP701A6
CYP701A9
CYP701A8
Sister lineage
Pharoideae
BOP
Bambusoideae
Oryzoideae
Pooideae
Core grass
PACMAD
Chloridoideae
Panicoideae
Pl06g08600 #17518
N3-N0
Pharus latifolius
Bradi1g37547 #4823
N3-N1
Brachypodium distachyon
N3-N2
TraesCS7B01G265800 #34373
Triticum aestivum B
eh_chr6.1096 #24364
N3-N4
Echinochloa haploclada
CsA302831.1 #12549
N3-N5
Cleistogenes songorica (A)
Oryza sativa

Figure S1

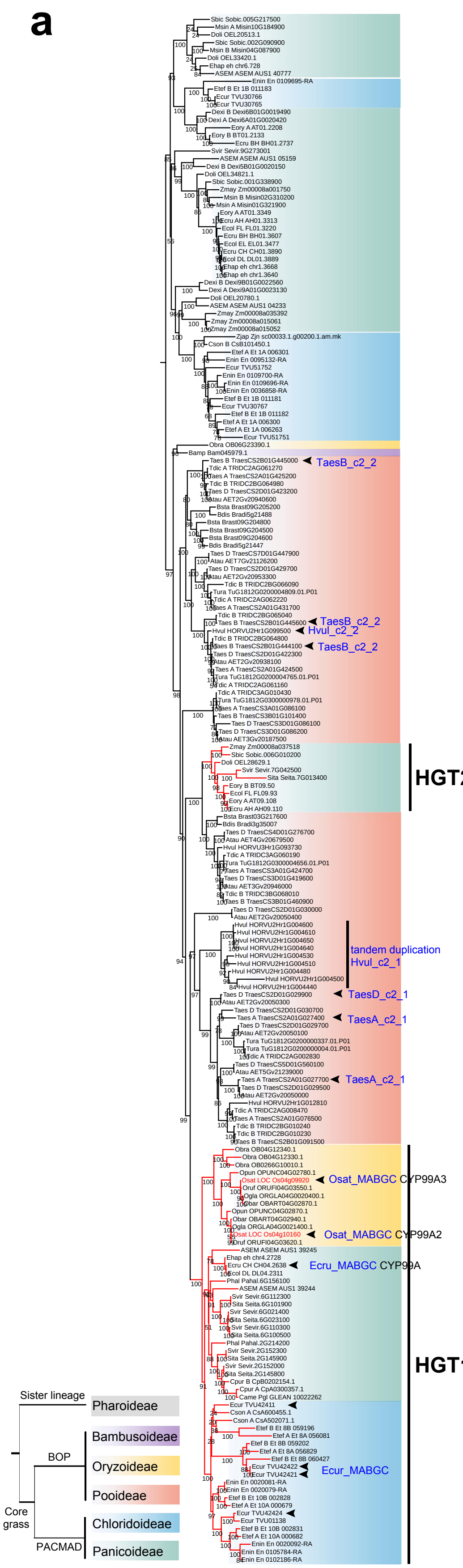

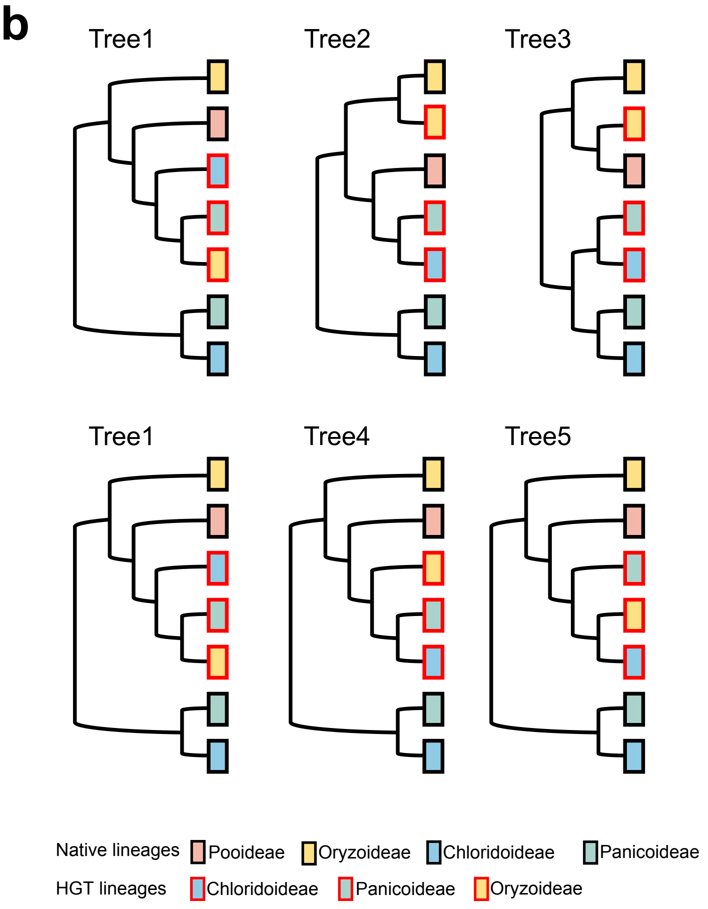

Test 1 : HGT or non-HGT

| Tree | bp-RELL | p-KH | p-SH | c-ELW |
|---|---|---|---|---|
| 1 | 1.0000 | 1.0000 | 1.0000 | 1.0000 |
| 2 | 0.0000- | 0.0000- | 0.0000- | 0.0000- |
| 3 | 0.0000- | 0.0000- | 0.0000- | 0.0000- |

Test 2 : relationship among PACMAD and *Oryza* HGT lineages

| Tree | bp-RELL | p-KH | p-SH | c-ELW |
|---|---|---|---|---|
| 1 | 0.2370 | 0.3870 | 0.5960 | 0.2367 |
| 4 | 0.3090 | 0.4050 | 0.5140 | 0.3094 |
| 5 | 0.4540 | 0.6130 | 1.0000 | 0.4538 |

Figure S2

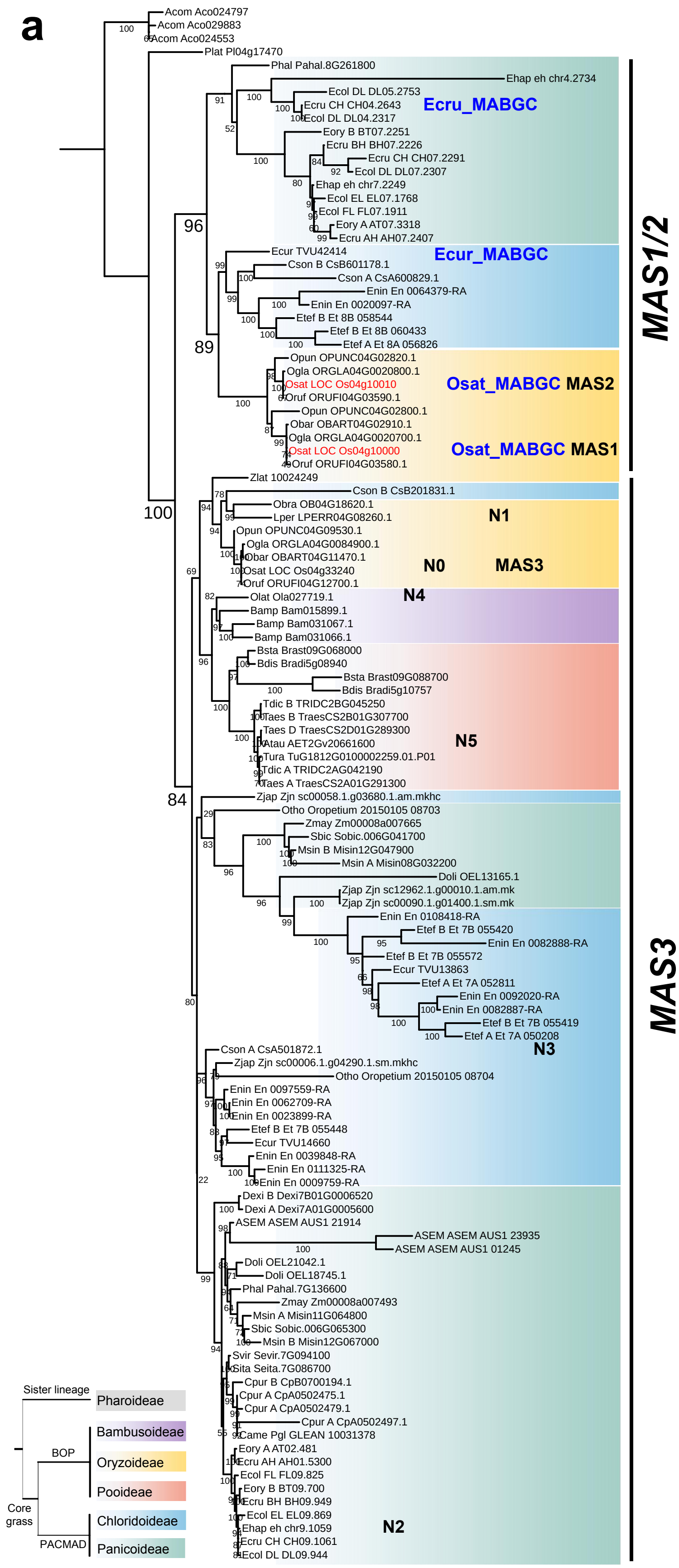

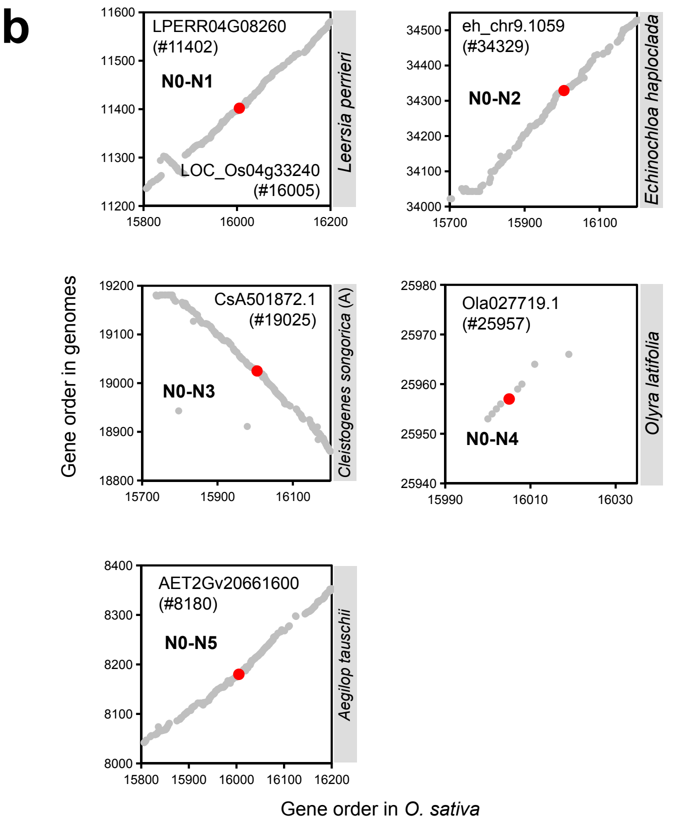

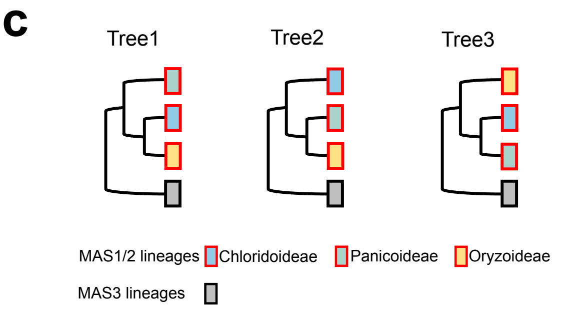

Test: relationship among PACMAD and *Oryza MAS1/2* lineages

| Tree | bp-RELL | p-KH | p-SH | c-ELW |
|---|---|---|---|---|
| 1 | 0.2580 | 0.2640 | 0.3110 | 0.2573 |
| 2 | 0.7110 | 0.7360 | 1.0000 | 0.7031 |
| 3 | 0.0310 - | 0.1040 | 0.2140 | 0.0395 - |

Figure S3

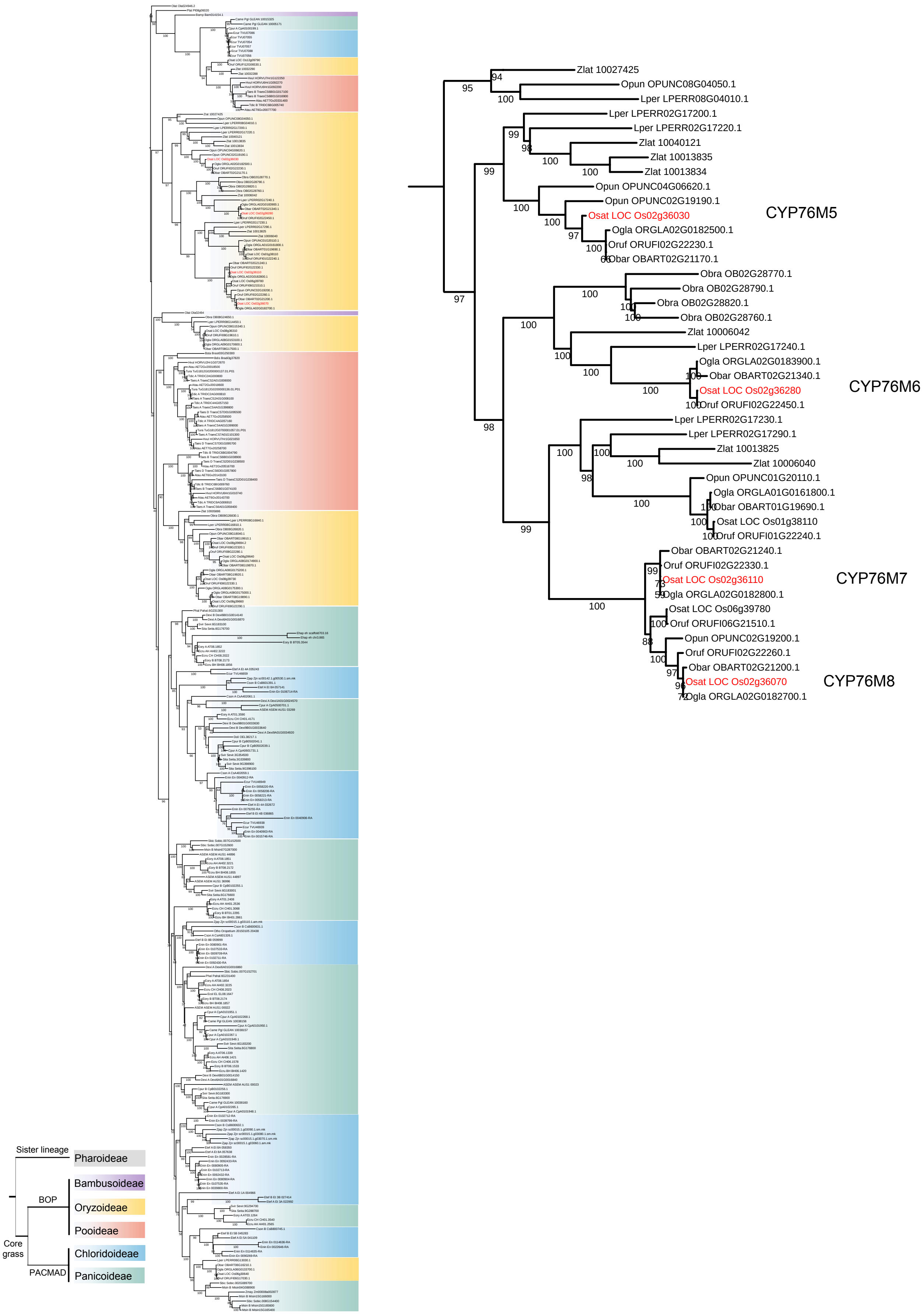
Zlat 10027425
Opun OPUNC08G04050.1
Lper LPERR08G04010.1
Lper LPERR02G17200.1
Lper LPERR02G17220.1
Zlat 10040121
Zlat 10013835
Zlat 10013834
Opun OPUNC04G06620.1
Opun OPUNC02G19190.1
Osat LOC Os02g36030
Ogla ORGLA02G0182500.1
Oruf ORUFI02G22230.1
Obar OBART02G21170.1
CYP76M5
Obra OB02G28770.1
Obra OB02G28790.1
Obra OB02G28820.1
Obra OB02G28760.1
Zlat 10006042
Lper LPERR02G17240.1
Ogla ORGLA02G0183900.1
Obar OBART02G21340.1
Osat LOC Os02g36280
Oruf ORUFI02G22450.1
CYP76M6
Lper LPERR02G17230.1
Lper LPERR02G17290.1
Zlat 10013825
Zlat 10006040
Opun OPUNC01G20110.1
Ogla ORGLA01G0161800.1
Obar OBART01G19690.1
Osat LOC Os01g38110
Oruf ORUFI01G22240.1
Obar OBART02G21240.1
Oruf ORUFI02G22330.1
Osat LOC Os02g36110
Ogla ORGLA02G0182800.1
CYP76M7
Osat LOC Os06g39780
Oruf ORUFI06G21510.1
Opun OPUNC02G19200.1
Oruf ORUFI02G22260.1
Obar OBART02G21200.1
Osat LOC Os02g36070
Ogla ORGLA02G0182700.1
CYP76M8
Sister lineage
Pharoideae
BOP
Bambusoideae
Oryzoideae
Pooideae
Core grass
PACMAD
Chloridoideae
Panicoideae

Figure S4

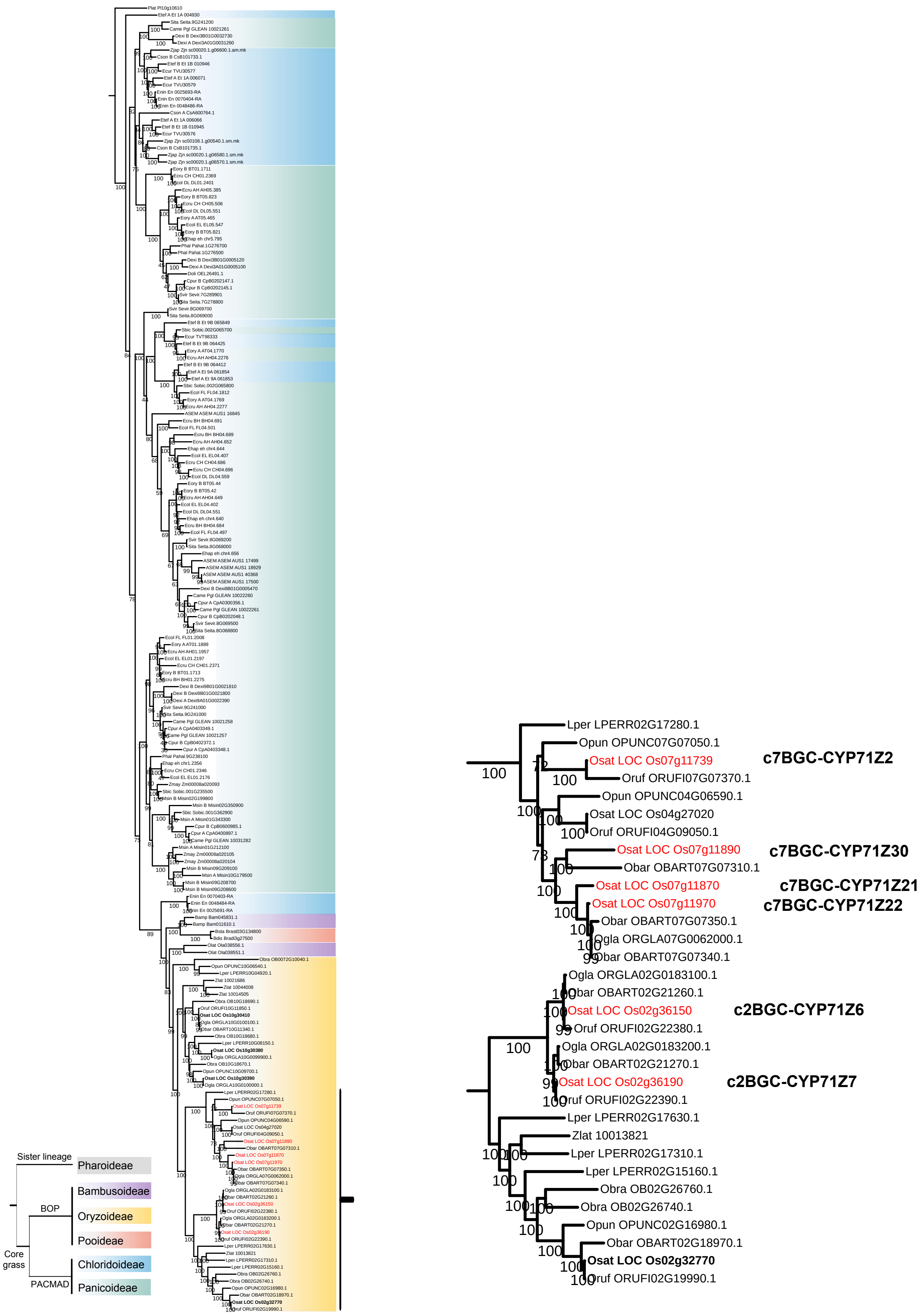
Sister lineage
Pharoideae
BOP
Bambusoideae
Oryzoideae
Pooideae
Core grass
PACMAD
Chloridoideae
Panicoideae
Lper LPERR02G17280.1
Opun OPUNC07G07050.1
Osat LOC Os07g11739
c7BGC-CYP71Z2
Oruf ORUFI07G07370.1
Opun OPUNC04G06590.1
Osat LOC Os04g27020
Oruf ORUFI04G09050.1
Osat LOC Os07g11890
c7BGC-CYP71Z30
Obar OBART07G07310.1
Osat LOC Os07g11870
c7BGC-CYP71Z21
Osat LOC Os07g11970
c7BGC-CYP71Z22
Obar OBART07G07350.1
Ogla ORGLA07G0062000.1
Obar OBART07G07340.1
Ogla ORGLA02G0183100.1
Obar OBART02G21260.1
Osat LOC Os02g36150
c2BGC-CYP71Z6
Oruf ORUFI02G22380.1
Ogla ORGLA02G0183200.1
Obar OBART02G21270.1
Osat LOC Os02g36190
c2BGC-CYP71Z7
Oruf ORUFI02G22390.1
Lper LPERR02G17630.1
Zlat 10013821
Lper LPERR02G17310.1
Lper LPERR02G15160.1
Obra OB02G26760.1
Obra OB02G26740.1
Opun OPUNC02G16980.1
Obar OBART02G18970.1
Osat LOC Os02g32770
Oruf ORUFI02G19990.1

Figure S5

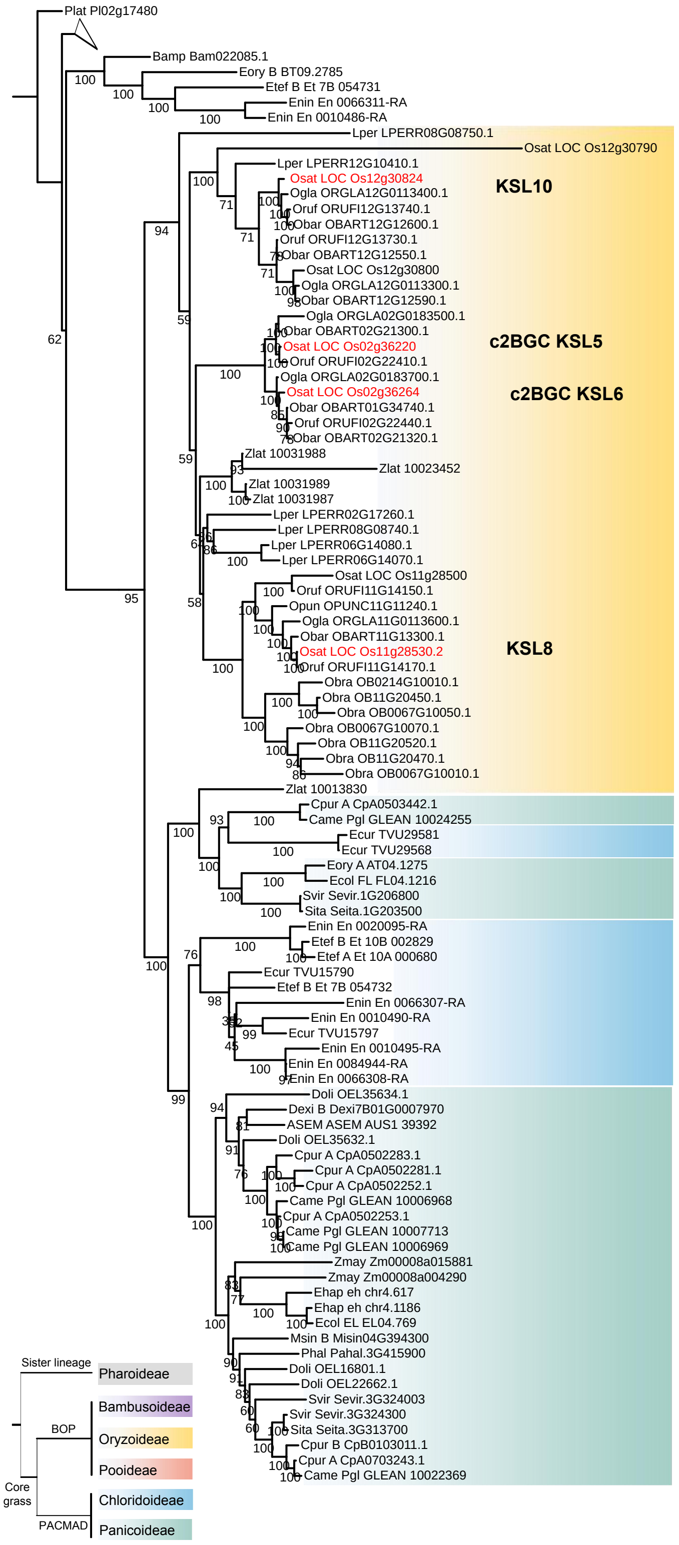

Plat Pl02g17480
Bamp Bam022085.1
Eory B BT09.2785
Etef B Et 7B 054731
Enin En 0066311-RA
Enin En 0010486-RA
Lper LPERR08G08750.1
Osat LOC Os12g30790
Lper LPERR12G10410.1
Osat LOC Os12g30824
Ogla ORGLA12G0113400.1
Oruf ORUFI12G13740.1
Obar OBART12G12600.1
Oruf ORUFI12G13730.1
Obar OBART12G12550.1
Osat LOC Os12g30800
Ogla ORGLA12G0113300.1
Obar OBART12G12590.1
KSL10
Ogla ORGLA02G0183500.1
Obar OBART02G21300.1
Osat LOC Os02g36220
Oruf ORUFI02G22410.1
Ogla ORGLA02G0183700.1
Osat LOC Os02g36264
Obar OBART01G34740.1
Oruf ORUFI02G22440.1
Obar OBART02G21320.1
c2BGC KSL5
c2BGC KSL6
Zlat 10031988
Zlat 10023452
Zlat 10031989
Zlat 10031987
Lper LPERR02G17260.1
Lper LPERR08G08740.1
Lper LPERR06G14080.1
Lper LPERR06G14070.1
Osat LOC Os11g28500
Oruf ORUFI11G14150.1
Opun OPUNC11G11240.1
Ogla ORGLA11G0113600.1
Obar OBART11G13300.1
Osat LOC Os11g28530.2
Oruf ORUFI11G14170.1
Obra OB0214G10010.1
Obra OB11G20450.1
Obra OB0067G10050.1
Obra OB0067G10070.1
Obra OB11G20520.1
Obra OB11G20470.1
Obra OB0067G10010.1
KSL8
Zlat 10013830
Cpur A CpA0503442.1
Came Pgl GLEAN 10024255
Ecur TVU29581
Ecur TVU29568
Eory A AT04.1275
Ecol FL FL04.1216
Svir Sevir.1G206800
Sita Seita.1G203500
Enin En 0020095-RA
Etef B Et 10B 002829
Etef A Et 10A 000680
Ecur TVU15790
Etef B Et 7B 054732
Enin En 0066307-RA
Enin En 0010490-RA
Ecur TVU15797
Enin En 0010495-RA
Enin En 0084944-RA
Enin En 0066308-RA
Doli OEL35634.1
Dexi B Dexi7B01G0007970
ASEM ASEM AUS1 39392
Doli OEL35632.1
Cpur A CpA0502283.1
Cpur A CpA0502281.1
Cpur A CpA0502252.1
Came Pgl GLEAN 10006968
Cpur A CpA0502253.1
Came Pgl GLEAN 10007713
Came Pgl GLEAN 10006969
Zmay Zm00008a015881
Zmay Zm00008a004290
Ehap eh chr4.617
Ehap eh chr4.1186
Ecol EL EL04.769
Msin B Misin04G394300
Phal Pahal.3G415900
Doli OEL16801.1
Doli OEL22662.1
Svir Sevir.3G324003
Svir Sevir.3G324300
Sita Seita.3G313700
Cpur B CpB0103011.1
Cpur A CpA0703243.1
Came Pgl GLEAN 10022369
Sister lineage
Pharoideae
BOP
Bambusoideae
Oryzoideae
Pooideae
Core grass
PACMAD
Chloridoideae
Panicoideae

Figure S6

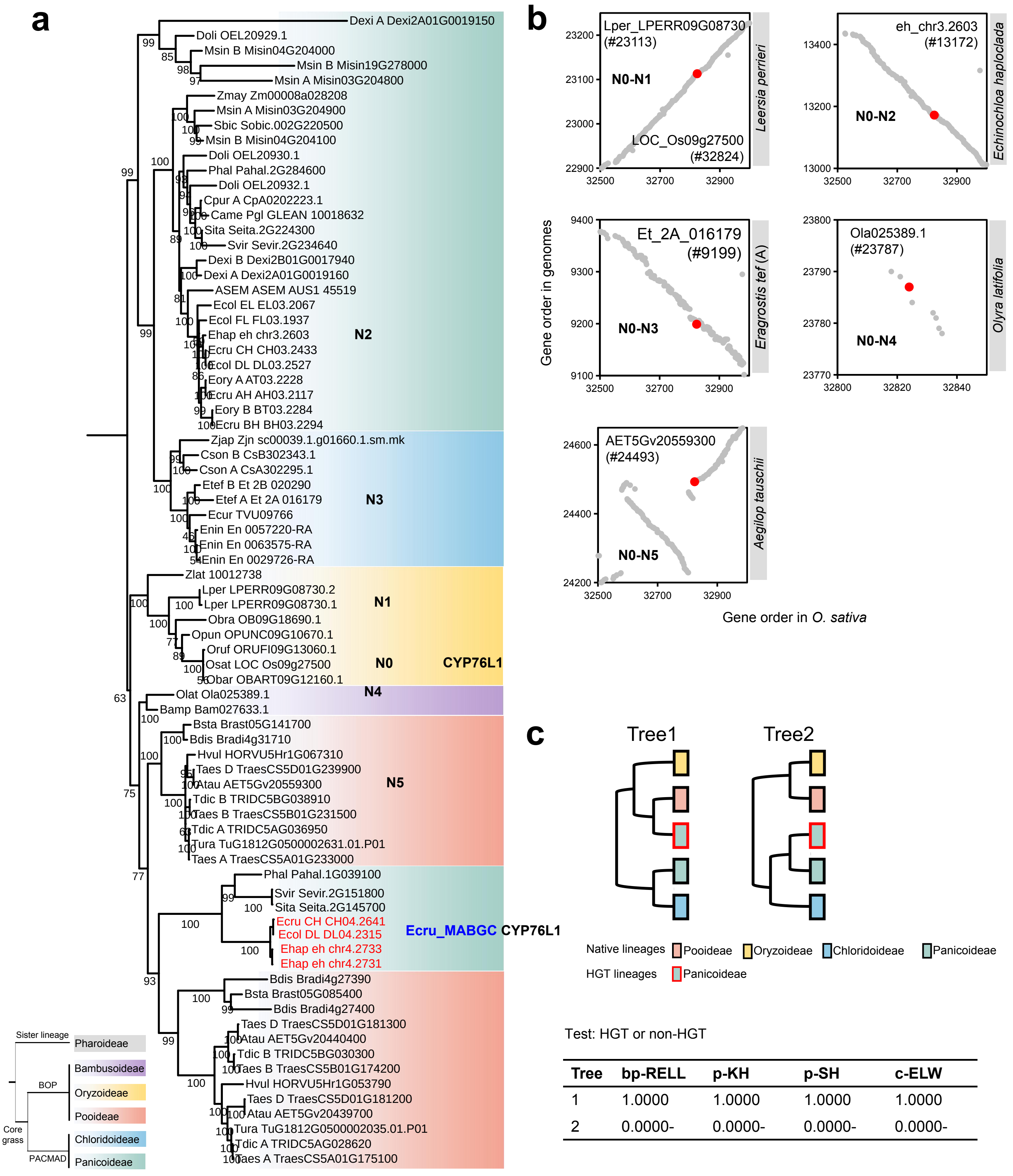

| Tree | bp-RELL | p-KH | p-SH | c-ELW |
|---|---|---|---|---|
| 1 | 1.0000 | 1.0000 | 1.0000 | 1.0000 |
| 2 | 0.0000- | 0.0000- | 0.0000- | 0.0000- |

Figure S7

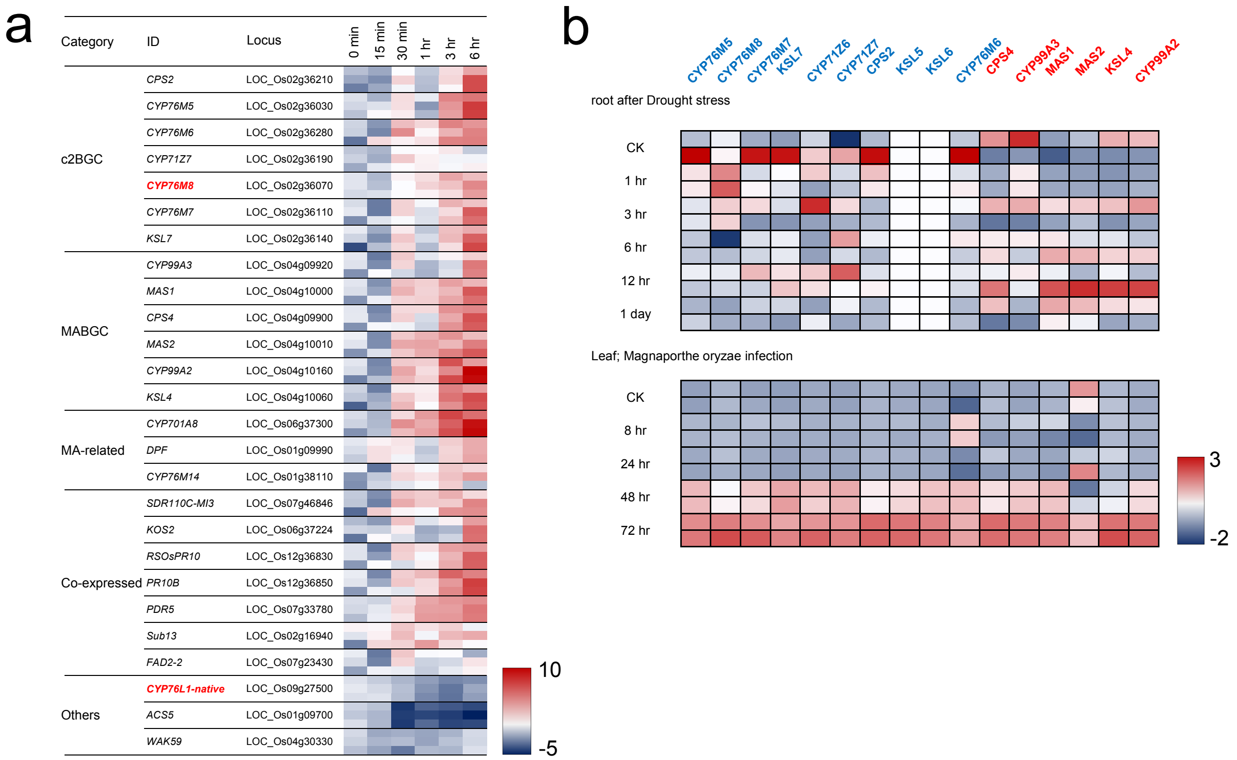
a
Category
ID
Locus
0 min
15 min
30 min
1 hr
3 hr
6 hr
c2BGC
CPS2 LOC_Os02g36210
CYP76M5 LOC_Os02g36030
CYP76M6 LOC_Os02g36280
CYP71Z7 LOC_Os02g36190
CYP76M8 LOC_Os02g36070
CYP76M7 LOC_Os02g36110
KSL7 LOC_Os02g36140
MABGC
CYP99A3 LOC_Os04g09920
MAS1 LOC_Os04g10000
CPS4 LOC_Os04g09900
MAS2 LOC_Os04g10010
CYP99A2 LOC_Os04g10160
KSL4 LOC_Os04g10060
MA-related
CYP701A8 LOC_Os06g37300
DPF LOC_Os01g09990
CYP76M14 LOC_Os01g38110
Co-expressed
SDR110C-MI3 LOC_Os07g46846
KOS2 LOC_Os06g37224
RSOsPR10 LOC_Os12g36830
PR10B LOC_Os12g36850
PDR5 LOC_Os07g33780
Sub13 LOC_Os02g16940
FAD2-2 LOC_Os07g23430
Others
CYP76L1-native LOC_Os09g27500
ACS5 LOC_Os01g09700
WAK59 LOC_Os04g30330
10
-5
b
CYP76M5 CYP76M8 CYP76M7 KSL7 CYP71Z6 CYP71Z7 CPS2 KSL5 KSL6 CYP76M6 CPS4 CYP99A3 MAS1 MAS2 KSL4 CYP99A2
root after Drought stress
CK
1 hr
3 hr
6 hr
12 hr
1 day
Leaf; Magnaporthe oryzae infection
CK
8 hr
24 hr
48 hr
72 hr
3
-2

### Figure S8

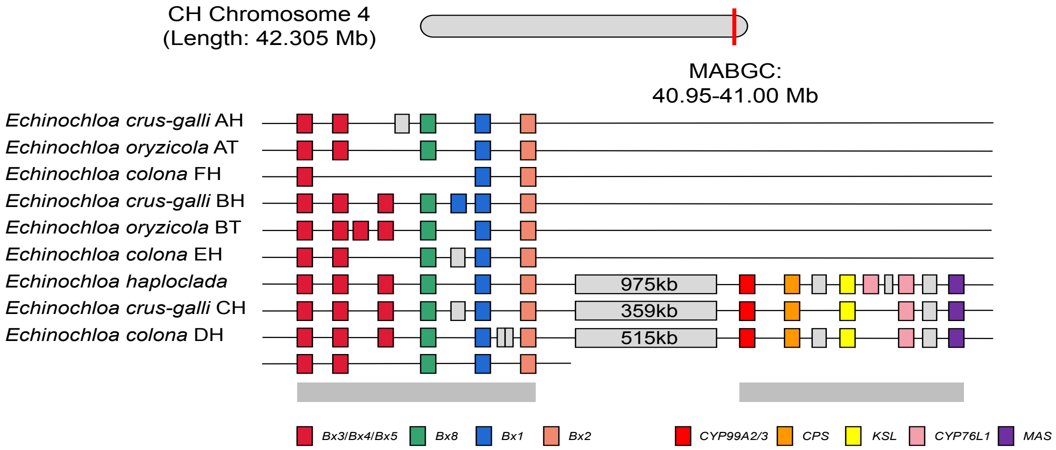
CH Chromosome 4
(Length: 42.305 Mb)
MABGC:
40.95-41.00 Mb
Echinochloa crus-galli AH
Echinochloa oryzicola AT
Echinochloa colona FH
Echinochloa crus-galli BH
Echinochloa oryzicola BT
Echinochloa colona EH
Echinochloa haploclada
Echinochloa crus-galli CH
Echinochloa colona DH
975kb
359kb
515kb
Bx3/Bx4/Bx5
Bx8
Bx1
Bx2
CYP99A2/3
CPS
KSL
CYP76L1
MAS

Figure S9

Table S1. A list of plant genomes used in this study

| Lineage | Sub-family | Species | Ploid | Version | Genome Abbr. |
|---|---|---|---|---|---|
| Outgroup | Bromeliaceae | *Ananas comosus* | 2 | F153 | Acom |
| Pharoideae | Pharoideae | *Pharus latifolius* | 2 | 60161 | Plat |
| BOP | Bambusoideae | *Olyra latifolia* | 2 | / | Olat |
| | | *Bonia amplexicaulis* | 6 | / | Bamp |
| | Oryzoideae | *Leersia perrieri* | 2 | GCA_000325765.[illegible] | Lper |
| | | *Oryza barthii* | 2 | GCA_000182155.[illegible] | Obar |
| | | *Oryza brachyantha* | 2 | GCA_000231095.[illegible] | Obra |
| | | *Oryza glaberrima* | 2 | GCA_000576495. | Ogla |
| | | *Oryza punctata* | 2 | GCA_000573905. | Opun |
| | | *Oryza rufipogon* | 2 | GCA_000817225. | Oruf |
| | | *Oryza sativa* | 2 | v7.0 | Osat |
| | | *Zizania latifolia* | 2 | v01 | Zlat |
| | Pooideae | *Aegilop tauschii* | 2 | v4.0 | Atau |
| | | *Brachypodium distachyon* | 2 | v3.1 | Bdis |
| | | *Brachypodium stacei* | 2 | v1.1 | Bsta |
| | | *Hordeum vulgare* | 2 | / | Hvul |
| | | *Triticum aestivum* | 6 | iwgsc_refseqv1.0 | Taes_A; Taes_B; Taes_D |
| | | *Triticum dicoccoides* | 4 | WEWSeq_v.1.0 | Tdic_A; Tdic_B |
| | | *Triticum urartu* | 2 | Tu | Tura |
| PACMAD | Panicoideae | *Alloteropsis semialata* | 2 | AUS1_V1.0 | Asem |
| | | *Cenchrus americanus* | 2 | Camericanus | Came |
| | | *Cenchrus purpureus* | 4 | GWHAORA00000 | Cpur_A; Cpur_B |
| | | *Dichanthelium oligosanth* | 2 | / | Doli |
| | | *Digitaria exilis* | 4 | CM05836 | Dexi_A; Dexi_B |
| | | *Miscanthus sinensis* | 4 | v7.1 | Msin_A; Msin_B |
| | | *Panicum hallii* | 2 | v3.1 | Phal |
| | | *Setaria italica* | 2 | v2.2 | Sita |
| | | *Setaria viridis* | 2 | v2.1 | Svir |
| | | *Sorghum bicolor* | 2 | v3.1.1 | Sbic |
| | | *Zea mays* | 2 | PH207_v1.1 | Zmay |
| | | *Echinochloa haploclada* | 2 | v1 | Ehap |
| | | *Echinochloa oryzicola* | 4 | v2 | Eory_AT; Eory_BT |
| | | *Echinochloa crus-galli* | 6 | v3 | Ecru_AH; Ecru_BH; Ecru_CH |
| | | *Echibochloa colona* | 6 | v1 | Ecol_DL; Ecol_EL; Ecol_FL |
| | Chloridoideae | *Cleistogenes songorica* | 4 | GWHANUQ00000 | Cson_A; Cson_B |
| | | *Eragrostis curvula* | 2 | / | Ecur |
| | | *Eragrostis tef* | 4 | v3.1 | Etef_A; Etef_B |
| | | *Eragrostis nindensis* | 4 | v2.1 | Enin |
| | | *Oropetium thomaeum* | 2 | Othomaeum | Otho |
| | | *Zoysia japonica* | 4 | Zjaponica_r1.1 | Zjap |

Table S2. Homologs of key genes (CPS4/KSL4/CYP99A/CYP76L/MAS) in momilactone biosynthetic gene cluster in grass

| Bait_Gene | Sub-family | Species | Gene_ID | Cluster/Pair | Cluster_Name | Gene_Name | Note |
|---|---|---|---|---|---|---|---|
| CPS4 | Pooideae | Atau | AET2Gv20049600 | clustered | Atau_c2_1 | | |
| KSL4 | Pooideae | Atau | AET2Gv20049900 | clustered | Atau_c2_1 | | |
| CYP99A2/3 | Pooideae | Atau | AET2Gv20050000 | clustered | Atau_c2_1 | | |
| CYP99A2/3 | Pooideae | Atau | AET2Gv20050100 | clustered | Atau_c2_1 | | |
| CYP99A2/3 | Pooideae | Atau | AET2Gv20050300 | clustered | Atau_c2_1 | | |
| CYP99A2/3 | Pooideae | Atau | AET2Gv20050400 | clustered | Atau_c2_1 | | |
| KSL4 | Pooideae | Atau | AET2Gv20050500 | clustered | Atau_c2_1 | | |
| KSL4 | Pooideae | Atau | AET2Gv20050600 | clustered | Atau_c2_1 | | |
| CPS4 | Pooideae | Atau | AET2Gv20050700 | clustered | Atau_c2_1 | | |
| CYP99A2/3 | Pooideae | Atau | AET2Gv20938100 | clustered | Atau_c2_2 | | |
| KSL4 | Pooideae | Atau | AET2Gv20939600 | clustered | Atau_c2_2 | | |
| KSL4 | Pooideae | Atau | AET2Gv20939800 | clustered | Atau_c2_2 | | |
| KSL4 | Pooideae | Atau | AET2Gv20940100 | clustered | Atau_c2_2 | | |
| CYP99A2/3 | Pooideae | Atau | AET2Gv20940600 | clustered | Atau_c2_2 | | |
| CYP701A8 | Pooideae | Atau | AET2Gv20948700 | | | | |
| CYP99A2/3 | Pooideae | Atau | AET2Gv20953300 | | | | |
| CPS4 | Pooideae | Atau | AET2Gv21176900 | | | | |
| CYP99A2/3 | Pooideae | Atau | AET3Gv20187500 | | | | |
| CYP99A2/3 | Pooideae | Atau | AET3Gv20946000 | | | | |
| CYP99A2/3 | Pooideae | Atau | AET4Gv20679500 | | | | |
| CYP76L1 | Pooideae | Atau | AET5Gv20439700 | | | | |
| CYP76L1 | Pooideae | Atau | AET5Gv20440400 | | | | |
| CYP76L1 | Pooideae | Atau | AET5Gv20559300 | | | | |
| CYP99A2/3 | Pooideae | Atau | AET5Gv21239000 | | | | |
| CPS4 | Pooideae | Atau | AET6Gv20453800 | | | | |
| CYP701A8 | Pooideae | Atau | AET7Gv20894800 | | | | |
| CYP701A8 | Pooideae | Atau | AET7Gv20894900 | | | | |
| CYP99A2/3 | Pooideae | Atau | AET7Gv21126200 | | | | |
| CYP701A8 | Pooideae | Bdis | Bradi1g37547 | | | | |
| CYP701A8 | Pooideae | Bdis | Bradi1g37560 | | | | |
| CYP701A8 | Pooideae | Bdis | Bradi1g37576 | | | | |
| CYP99A2/3 | Pooideae | Bdis | Bradi3g35007 | | | | |
| CYP76L1 | Pooideae | Bdis | Bradi4g27390 | | | | |
| CYP76L1 | Pooideae | Bdis | Bradi4g27400 | | | | |
| CYP76L1 | Pooideae | Bdis | Bradi4g31710 | | | | |
| KSL4 | Pooideae | Bdis | Bradi5g21387 | clustered | Bdis_c2_2 | | |
| KSL4 | Pooideae | Bdis | Bradi5g21440 | clustered | Bdis_c2_2 | | |
| CYP99A2/3 | Pooideae | Bdis | Bradi5g21447 | clustered | Bdis_c2_2 | | |
| KSL4 | Pooideae | Bdis | Bradi5g21480 | clustered | Bdis_c2_2 | | |
| CYP99A2/3 | Pooideae | Bdis | Bradi5g21488 | clustered | Bdis_c2_2 | | |
| KSL4 | Pooideae | Bdis | Bradi5g21497 | clustered | Bdis_c2_2 | | |
| CYP99A2/3 | Pooideae | Bsta | Brast03G217600 | | | | |
| CYP76L1 | Pooideae | Bsta | Brast05G085400 | | | | |
| CYP76L1 | Pooideae | Bsta | Brast05G141700 | | | | |
| CYP701A8 | Pooideae | Bsta | Brast07G150200 | | | | |
| CYP701A8 | Pooideae | Bsta | Brast07G150400 | | | | |
| CYP701A8 | Pooideae | Bsta | Brast07G150700 | | | | |
| KSL4 | Pooideae | Bsta | Brast09G203900 | clustered | Bsta_c2_2 | | |
| KSL4 | Pooideae | Bsta | Brast09G204400 | clustered | Bsta_c2_2 | | |
| CYP99A2/3 | Pooideae | Bsta | Brast09G204500 | clustered | Bsta_c2_2 | | |
| CYP99A2/3 | Pooideae | Bsta | Brast09G204600 | clustered | Bsta_c2_2 | | |
| CYP99A2/3 | Pooideae | Bsta | Brast09G204800 | clustered | Bsta_c2_2 | | |
| KSL4 | Pooideae | Bsta | Brast09G205100 | clustered | Bsta_c2_2 | | |
| CYP99A2/3 | Pooideae | Bsta | Brast09G205200 | clustered | Bsta_c2_2 | | |
| CYP701A8 | Pooideae | Hvul | HORVU0Hr1G021760 | | | | |
| CYP99A2/3 | Pooideae | Hvul | HORVU2Hr1G004440 | clustered | Hvul_c2_1 | | |
| CYP99A2/3 | Pooideae | Hvul | HORVU2Hr1G004480 | clustered | Hvul_c2_1 | | |
| CYP99A2/3 | Pooideae | Hvul | HORVU2Hr1G004500 | clustered | Hvul_c2_1 | | |
| CYP99A2/3 | Pooideae | Hvul | HORVU2Hr1G004510 | clustered | Hvul_c2_1 | | |
| CYP99A2/3 | Pooideae | Hvul | HORVU2Hr1G004530 | clustered | Hvul_c2_1 | | |
| KSL4 | Pooideae | Hvul | HORVU2Hr1G004540 | clustered | Hvul_c2_1 | | |
| CYP99A2/3 | Pooideae | Hvul | HORVU2Hr1G004600 | clustered | Hvul_c2_1 | | |
| CYP99A2/3 | Pooideae | Hvul | HORVU2Hr1G004610 | clustered | Hvul_c2_1 | | |
| CPS4 | Pooideae | Hvul | HORVU2Hr1G004620 | clustered | Hvul_c2_1 | | |
| CYP99A2/3 | Pooideae | Hvul | HORVU2Hr1G004640 | clustered | Hvul_c2_1 | | |
| CYP99A2/3 | Pooideae | Hvul | HORVU2Hr1G004650 | clustered | Hvul_c2_1 | | |
| CYP99A2/3 | Pooideae | Hvul | HORVU2Hr1G012810 | | | | |
| KSL4 | Pooideae | Hvul | HORVU2Hr1G099440 | clustered | Hvul_c2_2 | | |
| KSL4 | Pooideae | Hvul | HORVU2Hr1G099480 | clustered | Hvul_c2_2 | | |
| CYP99A2/3 | Pooideae | Hvul | HORVU2Hr1G099500 | clustered | Hvul_c2_2 | | |
| CYP701A8 | Pooideae | Hvul | HORVU2Hr1G100300 | | | | |
| CYP99A2/3 | Pooideae | Hvul | HORVU3Hr1G093730 | | | | |
| CYP76L1 | Pooideae | Hvul | HORVU5Hr1G053790 | | | | |
| CYP76L1 | Pooideae | Hvul | HORVU5Hr1G067310 | | | | |
| KSL4 | Pooideae | Hvul | HORVU5Hr1G082380 | paired | Hvul_c5 | | |

| | | | | | |
|---|---|---|---|---|---|
| CPS4 | Pooideae | Hvul | HORVU5Hr1G082390 | paired | Hvul_c5 |
| CPS4 | Pooideae | Taes | TraesCS2A01G027100 | clustered | TaesA_c2_1 |
| CPS4 | Pooideae | Taes | TraesCS2A01G027200 | clustered | TaesA_c2_1 |
| CYP99A2/3 | Pooideae | Taes | TraesCS2A01G027400 | clustered | TaesA_c2_1 |
| KSL4 | Pooideae | Taes | TraesCS2A01G027600 | clustered | TaesA_c2_1 |
| CYP99A2/3 | Pooideae | Taes | TraesCS2A01G027700 | clustered | TaesA_c2_1 |
| CPS4 | Pooideae | Taes | TraesCS2A01G027800 | clustered | TaesA_c2_1 |
| KSL4 | Pooideae | Taes | TraesCS2A01G028100 | clustered | TaesA_c2_1 |
| CPS4 | Pooideae | Taes | TraesCS2A01G028200 | clustered | TaesA_c2_1 |
| CYP99A2/3 | Pooideae | Taes | TraesCS2A01G076500 | | |
| CYP99A2/3 | Pooideae | Taes | TraesCS2A01G424500 | clustered | TaesA_c2_2 |
| KSL4 | Pooideae | Taes | TraesCS2A01G425100 | clustered | TaesA_c2_2 |
| CYP99A2/3 | Pooideae | Taes | TraesCS2A01G425200 | clustered | TaesA_c2_2 |
| KSL4 | Pooideae | Taes | TraesCS2A01G425400 | clustered | TaesA_c2_2 |
| CYP701A8 | Pooideae | Taes | TraesCS2A01G429600 | | |
| CYP99A2/3 | Pooideae | Taes | TraesCS2A01G431700 | | |
| CPS4 | Pooideae | Taes | TraesCS2A01G535300 | | |
| CPS4 | Pooideae | Taes | TraesCS2B01G040400 | clustered | TaesB_c2_1 |
| CPS4 | Pooideae | Taes | TraesCS2B01G040800 | clustered | TaesB_c2_1 |
| KSL4 | Pooideae | Taes | TraesCS2B01G040900 | clustered | TaesB_c2_1 |
| KSL4 | Pooideae | Taes | TraesCS2B01G041000 | clustered | TaesB_c2_1 |
| CPS4 | Pooideae | Taes | TraesCS2B01G041100 | clustered | TaesB_c2_1 |
| CYP99A2/3 | Pooideae | Taes | TraesCS2B01G091500 | | |
| CPS4 | Pooideae | Taes | TraesCS2B01G330000 | | |
| CYP99A2/3 | Pooideae | Taes | TraesCS2B01G444100 | clustered | TaesB_c2_2 |
| KSL4 | Pooideae | Taes | TraesCS2B01G444900 | clustered | TaesB_c2_2 |
| CYP99A2/3 | Pooideae | Taes | TraesCS2B01G445000 | clustered | TaesB_c2_2 |
| KSL4 | Pooideae | Taes | TraesCS2B01G445100 | clustered | TaesB_c2_2 |
| KSL4 | Pooideae | Taes | TraesCS2B01G445200 | clustered | TaesB_c2_2 |
| CYP701A8 | Pooideae | Taes | TraesCS2B01G445300 | | |
| CYP701A8 | Pooideae | Taes | TraesCS2B01G445400 | | |
| CPS4 | Pooideae | Taes | TraesCS2B01G445500 | clustered | TaesB_c2_2 |
| CYP99A2/3 | Pooideae | Taes | TraesCS2B01G445600 | clustered | TaesB_c2_2 |
| KSL4 | Pooideae | Taes | TraesCS2B01G445700 | clustered | TaesB_c2_2 |
| KSL4 | Pooideae | Taes | TraesCS2B01G450000 | paired | |
| CYP701A8 | Pooideae | Taes | TraesCS2B01G450100 | paired | |
| CYP701A8 | Pooideae | Taes | TraesCS2B01G450200 | paired | |
| KSL4 | Pooideae | Taes | TraesCS2B01G533600 | | |
| CPS4 | Pooideae | Taes | TraesCS2B01G565800 | | |
| CPS4 | Pooideae | Taes | TraesCS2B01G566300 | | |
| KSL4 | Pooideae | Taes | TraesCS2D01G029400 | clustered | TaesD_c2_1 |
| CYP99A2/3 | Pooideae | Taes | TraesCS2D01G029500 | clustered | TaesD_c2_1 |
| CPS4 | Pooideae | Taes | TraesCS2D01G029600 | clustered | TaesD_c2_1 |
| CYP99A2/3 | Pooideae | Taes | TraesCS2D01G029700 | clustered | TaesD_c2_1 |
| CYP99A2/3 | Pooideae | Taes | TraesCS2D01G029900 | clustered | TaesD_c2_1 |
| CYP99A2/3 | Pooideae | Taes | TraesCS2D01G030000 | clustered | TaesD_c2_1 |
| KSL4 | Pooideae | Taes | TraesCS2D01G030100 | clustered | TaesD_c2_1 |
| KSL4 | Pooideae | Taes | TraesCS2D01G030200 | clustered | TaesD_c2_1 |
| CPS4 | Pooideae | Taes | TraesCS2D01G030300 | clustered | TaesD_c2_1 |
| CYP99A2/3 | Pooideae | Taes | TraesCS2D01G030700 | clustered | TaesD_c2_1 |
| CYP99A2/3 | Pooideae | Taes | TraesCS2D01G422300 | clustered | TaesD_c2_2 |
| KSL4 | Pooideae | Taes | TraesCS2D01G423100 | clustered | TaesD_c2_2 |
| CYP99A2/3 | Pooideae | Taes | TraesCS2D01G423200 | clustered | TaesD_c2_2 |
| KSL4 | Pooideae | Taes | TraesCS2D01G423300 | clustered | TaesD_c2_2 |
| CYP701A8 | Pooideae | Taes | TraesCS2D01G427200 | | |
| CYP99A2/3 | Pooideae | Taes | TraesCS2D01G429700 | | |
| KSL4 | Pooideae | Taes | TraesCS2D01G506000 | | |
| CPS4 | Pooideae | Taes | TraesCS2D01G537200 | | |
| CYP99A2/3 | Pooideae | Taes | TraesCS3A01G086100 | | |
| CYP99A2/3 | Pooideae | Taes | TraesCS3A01G424700 | | |
| CYP99A2/3 | Pooideae | Taes | TraesCS3B01G101400 | | |
| CYP99A2/3 | Pooideae | Taes | TraesCS3B01G460900 | | |
| CYP99A2/3 | Pooideae | Taes | TraesCS3D01G086100 | | |
| CYP99A2/3 | Pooideae | Taes | TraesCS3D01G086200 | | |
| CYP99A2/3 | Pooideae | Taes | TraesCS3D01G419600 | | |
| CYP99A2/3 | Pooideae | Taes | TraesCS4D01G276700 | | |
| CYP76L1 | Pooideae | Taes | TraesCS5A01G175100 | | |
| CYP76L1 | Pooideae | Taes | TraesCS5A01G233000 | | |
| CYP76L1 | Pooideae | Taes | TraesCS5B01G174200 | | |
| CYP76L1 | Pooideae | Taes | TraesCS5B01G231500 | | |
| CYP76L1 | Pooideae | Taes | TraesCS5D01G181200 | | |
| CYP76L1 | Pooideae | Taes | TraesCS5D01G181300 | | |
| CYP76L1 | Pooideae | Taes | TraesCS5D01G239900 | | |
| CYP99A2/3 | Pooideae | Taes | TraesCS5D01G560100 | | |
| KSL4 | Pooideae | Taes | TraesCS6B01G034000 | | |
| CYP701A8 | Pooideae | Taes | TraesCS7A01G362200 | | |
| CYP701A8 | Pooideae | Taes | TraesCS7A01G362300 | | |

| | | | | | |
|---|---|---|---|---|---|
| CYP701A8 | Pooideae | Taes | TraesCS7B01G265800 | | |
| CPS4 | Pooideae | Taes | TraesCS7D01G097000 | | |
| CYP701A8 | Pooideae | Taes | TraesCS7D01G360700 | | |
| CYP99A2/3 | Pooideae | Taes | TraesCS7D01G447900 | | |
| CYP99A2/3 | Pooideae | Tdic | TRIDC2AG002830 | clustered | TdicA_c2_1 |
| CPS4 | Pooideae | Tdic | TRIDC2AG002840 | clustered | TdicA_c2_1 |
| CPS4 | Pooideae | Tdic | TRIDC2AG002850 | clustered | TdicA_c2_1 |
| CYP99A2/3 | Pooideae | Tdic | TRIDC2AG008470 | | |
| CYP99A2/3 | Pooideae | Tdic | TRIDC2AG061160 | clustered | TdicA_c2_2 |
| KSL4 | Pooideae | Tdic | TRIDC2AG061250 | clustered | TdicA_c2_2 |
| KSL4 | Pooideae | Tdic | TRIDC2AG061260 | clustered | TdicA_c2_2 |
| CYP99A2/3 | Pooideae | Tdic | TRIDC2AG061270 | clustered | TdicA_c2_2 |
| KSL4 | Pooideae | Tdic | TRIDC2AG061360 | clustered | TdicA_c2_2 |
| CYP701A8 | Pooideae | Tdic | TRIDC2AG061850 | | |
| CYP99A2/3 | Pooideae | Tdic | TRIDC2AG062220 | | |
| CPS4 | Pooideae | Tdic | TRIDC2BG003190 | paired | TdicB_c2_1 |
| KSL4 | Pooideae | Tdic | TRIDC2BG003240 | paired | TdicB_c2_1 |
| CYP99A2/3 | Pooideae | Tdic | TRIDC2BG010230 | | |
| CYP99A2/3 | Pooideae | Tdic | TRIDC2BG010240 | | |
| CYP99A2/3 | Pooideae | Tdic | TRIDC2BG064800 | | |
| KSL4 | Pooideae | Tdic | TRIDC2BG064970 | clustered | TdicB_c2_2 |
| CYP99A2/3 | Pooideae | Tdic | TRIDC2BG064980 | clustered | TdicB_c2_2 |
| KSL4 | Pooideae | Tdic | TRIDC2BG065000 | clustered | TdicB_c2_2 |
| KSL4 | Pooideae | Tdic | TRIDC2BG065010 | clustered | TdicB_c2_2 |
| CYP701A8 | Pooideae | Tdic | TRIDC2BG065020 | | |
| CYP701A8 | Pooideae | Tdic | TRIDC2BG065030 | | |
| CYP99A2/3 | Pooideae | Tdic | TRIDC2BG065040 | clustered | TdicB_c2_2 |
| KSL4 | Pooideae | Tdic | TRIDC2BG065050 | clustered | TdicB_c2_2 |
| CYP701A8 | Pooideae | Tdic | TRIDC2BG065680 | | |
| CYP701A8 | Pooideae | Tdic | TRIDC2BG065700 | | |
| CYP99A2/3 | Pooideae | Tdic | TRIDC2BG066090 | | |
| CYP99A2/3 | Pooideae | Tdic | TRIDC3AG010430 | | |
| CYP99A2/3 | Pooideae | Tdic | TRIDC3AG060190 | | |
| CYP99A2/3 | Pooideae | Tdic | TRIDC3BG068010 | | |
| CYP76L1 | Pooideae | Tdic | TRIDC5AG028620 | | |
| CYP76L1 | Pooideae | Tdic | TRIDC5AG036950 | | |
| CYP76L1 | Pooideae | Tdic | TRIDC5BG030300 | | |
| CYP76L1 | Pooideae | Tdic | TRIDC5BG038910 | | |
| KSL4 | Pooideae | Tdic | TRIDC6BG003980 | | |
| KSL4 | Pooideae | Tdic | TRIDC6BG003990 | | |
| CYP701A8 | Pooideae | Tdic | TRIDC7AG050600 | | |
| CYP701A8 | Pooideae | Tdic | TRIDC7AG050610 | | |
| CYP701A8 | Pooideae | Tdic | TRIDC7BG043260 | | |
| CYP99A2/3 | Pooideae | Tura | TuG1812G0200000004.01 | | |
| CPS4 | Pooideae | Tura | TuG1812G0200000312.01 | | |
| CYP99A2/3 | Pooideae | Tura | TuG1812G0200000337.01 | clustered | Tura_c2_1 |
| CPS4 | Pooideae | Tura | TuG1812G0200000339.01 | clustered | Tura_c2_1 |
| KSL4 | Pooideae | Tura | TuG1812G0200000340.01 | clustered | Tura_c2_1 |
| KSL4 | Pooideae | Tura | TuG1812G0200000341.01 | clustered | Tura_c2_1 |
| CPS4 | Pooideae | Tura | TuG1812G0200000342.01 | clustered | Tura_c2_1 |
| KSL4 | Pooideae | Tura | TuG1812G0200004755.01 | clustered | Tura_c2_2 |
| KSL4 | Pooideae | Tura | TuG1812G0200004756.01 | clustered | Tura_c2_2 |
| KSL4 | Pooideae | Tura | TuG1812G0200004757.01 | clustered | Tura_c2_2 |
| CYP99A2/3 | Pooideae | Tura | TuG1812G0200004765.01 | clustered | Tura_c2_2 |
| KSL4 | Pooideae | Tura | TuG1812G0200004769.01 | clustered | Tura_c2_2 |
| KSL4 | Pooideae | Tura | TuG1812G0200004779.01 | clustered | Tura_c2_2 |
| CYP99A2/3 | Pooideae | Tura | TuG1812G0200004809.01 | | |
| KSL4 | Pooideae | Tura | TuG1812G0200004823.01 | | |
| CYP701A8 | Pooideae | Tura | TuG1812G0200004824.01 | | |
| KSL4 | Pooideae | Tura | TuG1812G0200005492.01 | | |
| CPS4 | Pooideae | Tura | TuG1812G0200005796.01 | | |
| CYP99A2/3 | Pooideae | Tura | TuG1812G0300000978.01 | | |
| CYP99A2/3 | Pooideae | Tura | TuG1812G0300004656.01 | | |
| CYP76L1 | Pooideae | Tura | TuG1812G0500002035.01 | | |
| CYP76L1 | Pooideae | Tura | TuG1812G0500002631.01 | | |
| KSL4 | Pooideae | Tura | TuG1812G0600000233.01 | | |
| CYP701A8 | Pooideae | Tura | TuG1812G0700003978.01 | | |
| CYP701A8 | Pooideae | Tura | TuG1812G0700003979.01 | | |
| CPS4 | Pooideae | Tura | TuG1812U0000074100.01 | | |
| CYP99A2/3 | Panicoideae | ASEM | ASEM_AUS1_04233 | | |
| CYP99A2/3 | Panicoideae | ASEM | ASEM_AUS1_05159 | | |
| CPS4 | Panicoideae | ASEM | ASEM_AUS1_13415 | | |
| CPS4 | Panicoideae | ASEM | ASEM_AUS1_18850 | | |
| CPS4 | Panicoideae | ASEM | ASEM_AUS1_18852 | | |
| CYP701A8 | Panicoideae | ASEM | ASEM_AUS1_23329 | | |
| KSL4 | Panicoideae | ASEM | ASEM_AUS1_30767 | | |
| CYP99A2/3 | Panicoideae | ASEM | ASEM_AUS1_39244 | | |

| | | | | | | |
|---|---|---|---|---|---|---|
| CYP99A2/3 | Panicoideae | ASEM | ASEM_AUS1_39245 | | | |
| CYP99A2/3 | Panicoideae | ASEM | ASEM_AUS1_40777 | | | |
| CYP76L1 | Panicoideae | ASEM | ASEM_AUS1_45519 | | | |
| CPS4 | Panicoideae | Came | Pgl_GLEAN_10011293 | | | |
| KSL4 | Panicoideae | Came | Pgl_GLEAN_10012752 | | | |
| CYP76L1 | Panicoideae | Came | Pgl_GLEAN_10018632 | | | |
| CYP99A2/3 | Panicoideae | Came | Pgl_GLEAN_10022262 | clustered | | |
| CPS4 | Panicoideae | Came | Pgl_GLEAN_10022268 | clustered | | |
| CPS4 | Panicoideae | Came | Pgl_GLEAN_10022271 | clustered | | |
| CPS4 | Panicoideae | Came | Pgl_GLEAN_10022272 | clustered | | |
| CYP701A8 | Panicoideae | Came | Pgl_GLEAN_10024915 | | | |
| CYP701A8 | Panicoideae | Came | Pgl_GLEAN_10038310 | | | |
| CYP701A8 | Panicoideae | Cpur | CpA0105446.1 | | | |
| CYP701A8 | Panicoideae | Cpur | CpA0105466.1 | | | |
| CYP76L1 | Panicoideae | Cpur | CpA0202223.1 | | | |
| CYP99A2/3 | Panicoideae | Cpur | CpA0300357.1 | paired | | |
| CPS4 | Panicoideae | Cpur | CpA0300359.1 | paired | | |
| CPS4 | Panicoideae | Cpur | CpA0300461.1 | | | |
| KSL4 | Panicoideae | Cpur | CpA0500792.1 | | | |
| KSL4 | Panicoideae | Cpur | CpA0500839.1 | | | |
| CPS4 | Panicoideae | Cpur | CpA0503468.1 | | | |
| CPS4 | Panicoideae | Cpur | CpA0503470.1 | | | |
| CYP701A8 | Panicoideae | Cpur | CpA0603606.1 | | | |
| CYP99A2/3 | Panicoideae | Cpur | CpB0202154.1 | paired | | |
| CPS4 | Panicoideae | Cpur | CpB0202155.1 | paired | | |
| CYP701A8 | Panicoideae | Cpur | CpB0502792.1 | | | |
| CPS4 | Panicoideae | Cpur | CpB0601648.1 | | | |
| CPS4 | Panicoideae | Cpur | CpB0601649.1 | | | |
| KSL4 | Panicoideae | Cpur | CpB0701686.1 | | | |
| KSL4 | Panicoideae | Cpur | CpB0701719.1 | | | |
| CPS4 | Panicoideae | Dexi | Dexi1A01G0017880 | | | |
| CPS4 | Panicoideae | Dexi | Dexi1B01G0016720 | | | |
| CPS4 | Panicoideae | Dexi | Dexi1B01G0016730 | | | |
| CYP76L1 | Panicoideae | Dexi | Dexi2A01G0019150 | | | |
| CYP76L1 | Panicoideae | Dexi | Dexi2A01G0019160 | | | |
| CYP76L1 | Panicoideae | Dexi | Dexi2B01G0017940 | | | |
| CYP701A8 | Panicoideae | Dexi | Dexi4A01G0015790 | | | |
| CYP701A8 | Panicoideae | Dexi | Dexi4A01G0015800 | | | |
| CPS4 | Panicoideae | Dexi | Dexi5A01G0018150 | | | |
| CPS4 | Panicoideae | Dexi | Dexi5B01G0018500 | | | |
| CYP99A2/3 | Panicoideae | Dexi | Dexi5B01G0020150 | | | |
| CYP99A2/3 | Panicoideae | Dexi | Dexi6A01G0020420 | | | |
| CYP99A2/3 | Panicoideae | Dexi | Dexi6B01G0019490 | | | |
| KSL4 | Panicoideae | Dexi | Dexi7A01G0018450 | | | |
| KSL4 | Panicoideae | Dexi | Dexi7B01G0019210 | | | |
| CYP99A2/3 | Panicoideae | Dexi | Dexi9A01G0023130 | | | |
| CYP99A2/3 | Panicoideae | Dexi | Dexi9B01G0022560 | | | |
| CYP701A8 | Panicoideae | Dexi | DexiUA01G0007890 | | | |
| CYP701A8 | Panicoideae | Dexi | DexiUA01G0007900 | | | |
| CYP99A2/3 | Panicoideae | Doli | OEL20513.1 | | | |
| CYP99A2/3 | Panicoideae | Doli | OEL20780.1 | | | |
| CYP76L1 | Panicoideae | Doli | OEL20929.1 | | | |
| CYP76L1 | Panicoideae | Doli | OEL20930.1 | | | |
| CYP76L1 | Panicoideae | Doli | OEL20932.1 | | | |
| CPS4 | Panicoideae | Doli | OEL23223.1 | | | |
| CYP701A8 | Panicoideae | Doli | OEL24076.1 | | | |
| CYP99A2/3 | Panicoideae | Doli | OEL28629.1 | | | |
| CYP99A2/3 | Panicoideae | Doli | OEL33420.1 | | | |
| CYP99A2/3 | Panicoideae | Doli | OEL34821.1 | | | |
| CYP99A2/3 | Panicoideae | Ecol | DL01.3889 | | | |
| CYP76L1 | Panicoideae | Ecol | DL03.2527 | | | |
| CYP99A2/3 | Panicoideae | Ecol | DL04.2311 | clustered | Ecol_MABGC | MABGC |
| CPS4 | Panicoideae | Ecol | DL04.2312 | clustered | Ecol_MABGC | MABGC |
| KSL4 | Panicoideae | Ecol | DL04.2314 | clustered | Ecol_MABGC | MABGC |
| CYP76L1 | Panicoideae | Ecol | DL04.2315 | clustered | Ecol_MABGC | MABGC |
| MAS1/2 | Panicoideae | Ecol | DL04.2317 | clustered | Ecol_MABGC | MABGC |
| MAS1/2 | Panicoideae | Ecol | DL05.2753 | | | |
| CYP701A8 | Panicoideae | Ecol | DL06.1065 | | | |
| CPS4 | Panicoideae | Ecol | DL07.2200 | | | |
| CPS4 | Panicoideae | Ecol | DL07.2201 | | | |
| MAS1/2 | Panicoideae | Ecol | DL07.2307 | | | |
| KSL4 | Panicoideae | Ecol | DL09.2533 | | | |
| CYP99A2/3 | Panicoideae | Ecol | EL01.3477 | | | |
| CYP76L1 | Panicoideae | Ecol | EL03.2067 | | | |
| CYP701A8 | Panicoideae | Ecol | EL06.934 | | | |
| CPS4 | Panicoideae | Ecol | EL07.1650 | | | |
| MAS1/2 | Panicoideae | Ecol | EL07.1768 | | | |

| | | | | | | |
|---|---|---|---|---|---|---|
| KSL4 | Panicoideae | Ecol | EL09.2272 | | | |
| CYP99A2/3 | Panicoideae | Ecol | FL01.3220 | | | |
| CYP76L1 | Panicoideae | Ecol | FL03.1937 | | | |
| CPS4 | Panicoideae | Ecol | FL04.1217 | | | |
| CYP701A8 | Panicoideae | Ecol | FL06.2082 | | | |
| MAS1/2 | Panicoideae | Ecol | FL07.1911 | | | |
| KSL4 | Panicoideae | Ecol | FL09.2178 | | | |
| CYP99A2/3 | Panicoideae | Ecol | FL09.93 | | | |
| CYP99A2/3 | Panicoideae | Ecru | AH01.3313 | | | |
| CYP76L1 | Panicoideae | Ecru | AH03.2117 | | | |
| CPS4 | Panicoideae | Ecru | AH04.1478 | | | |
| CYP701A8 | Panicoideae | Ecru | AH06.2079 | | | |
| MAS1/2 | Panicoideae | Ecru | AH07.2407 | | | |
| CYP99A2/3 | Panicoideae | Ecru | AH09.110 | | | |
| KSL4 | Panicoideae | Ecru | AH09.2264 | | | |
| CYP99A2/3 | Panicoideae | Ecru | BH01.2737 | | | |
| CYP99A2/3 | Panicoideae | Ecru | BH01.3607 | | | |
| CYP76L1 | Panicoideae | Ecru | BH03.2294 | | | |
| CPS4 | Panicoideae | Ecru | BH04.1488 | | | |
| CYP701A8 | Panicoideae | Ecru | BH06.1921 | | | |
| MAS1/2 | Panicoideae | Ecru | BH07.2226 | | | |
| CPS4 | Panicoideae | Ecru | BH07.3937 | | | |
| CPS4 | Panicoideae | Ecru | BH07.3939 | | | |
| KSL4 | Panicoideae | Ecru | BH09.2446 | | | |
| CYP99A2/3 | Panicoideae | Ecru | CH01.3890 | | | |
| CYP76L1 | Panicoideae | Ecru | CH03.2433 | | | |
| CYP99A2/3 | Panicoideae | Ecru | CH04.2638 | clustered | Ecru_MABGC | MABGC |
| CPS4 | Panicoideae | Ecru | CH04.2639 | clustered | Ecru_MABGC | MABGC |
| KSL4 | Panicoideae | Ecru | CH04.2640 | clustered | Ecru_MABGC | MABGC |
| CYP76L1 | Panicoideae | Ecru | CH04.2641 | clustered | Ecru_MABGC | MABGC |
| MAS1/2 | Panicoideae | Ecru | CH04.2643 | clustered | Ecru_MABGC | MABGC |
| CPS4 | Panicoideae | Ecru | CH07.2180 | | | |
| MAS1/2 | Panicoideae | Ecru | CH07.2291 | | | |
| KSL4 | Panicoideae | Ecru | CH09.2590 | | | |
| CYP99A2/3 | Panicoideae | Ehap | eh_chr1.3640 | | | |
| CYP99A2/3 | Panicoideae | Ehap | eh_chr1.3668 | | | |
| CYP76L1 | Panicoideae | Ehap | eh_chr3.2603 | | | |
| CYP99A2/3 | Panicoideae | Ehap | eh_chr4.2728 | clustered | Ehap_MABGC | MABGC |
| CPS4 | Panicoideae | Ehap | eh_chr4.2729 | clustered | Ehap_MABGC | MABGC |
| KSL4 | Panicoideae | Ehap | eh_chr4.27305 | clustered | Ehap_MABGC | MABGC |
| CYP76L1 | Panicoideae | Ehap | eh_chr4.2731 | clustered | Ehap_MABGC | MABGC |
| CYP76L1 | Panicoideae | Ehap | eh_chr4.2733 | clustered | Ehap_MABGC | MABGC |
| MAS1/2 | Panicoideae | Ehap | eh_chr4.2734 | clustered | Ehap_MABGC | MABGC |
| CYP701A8 | Panicoideae | Ehap | eh_chr6.1096 | | | |
| CYP99A2/3 | Panicoideae | Ehap | eh_chr6.728 | | | |
| CPS4 | Panicoideae | Ehap | eh_chr7.2132 | | | |
| MAS1/2 | Panicoideae | Ehap | eh_chr7.2249 | | | |
| KSL4 | Panicoideae | Ehap | eh_chr9.2617 | | | |
| CYP99A2/3 | Panicoideae | Eory | AT01.2208 | | | |
| CYP99A2/3 | Panicoideae | Eory | AT01.3349 | | | |
| CYP76L1 | Panicoideae | Eory | AT03.2228 | | | |
| CPS4 | Panicoideae | Eory | AT04.1277 | | | |
| CYP701A8 | Panicoideae | Eory | AT06.1831 | | | |
| MAS1/2 | Panicoideae | Eory | AT07.3318 | | | |
| CYP99A2/3 | Panicoideae | Eory | AT09.108 | | | |
| KSL4 | Panicoideae | Eory | AT09.2337 | | | |
| CYP99A2/3 | Panicoideae | Eory | BT01.2133 | | | |
| CYP76L1 | Panicoideae | Eory | BT03.2284 | | | |
| CPS4 | Panicoideae | Eory | BT04.1287 | | | |
| CYP701A8 | Panicoideae | Eory | BT06.2057 | | | |
| CPS4 | Panicoideae | Eory | BT07.2142 | | | |
| MAS1/2 | Panicoideae | Eory | BT07.2251 | | | |
| KSL4 | Panicoideae | Eory | BT09.2778 | | | |
| CYP99A2/3 | Panicoideae | Eory | BT09.50 | | | |
| CYP99A2/3 | Panicoideae | Msin | Misin01G321900 | | | |
| CYP99A2/3 | Panicoideae | Msin | Misin02G310200 | | | |
| CYP76L1 | Panicoideae | Msin | Misin03G204800 | | | |
| CYP76L1 | Panicoideae | Msin | Misin03G204900 | | | |
| CYP99A2/3 | Panicoideae | Msin | Misin04G087900 | | | |
| CYP76L1 | Panicoideae | Msin | Misin04G204000 | | | |
| CYP76L1 | Panicoideae | Msin | Misin04G204100 | | | |
| CPS4 | Panicoideae | Msin | Misin06G310100 | | | |
| CYP99A2/3 | Panicoideae | Msin | Misin10G184900 | clustered | | |
| CPS4 | Panicoideae | Msin | Misin10G188900 | clustered | | |
| CPS4 | Panicoideae | Msin | Misin10G189000 | clustered | | |
| KSL4 | Panicoideae | Msin | Misin11G207700 | | | |
| KSL4 | Panicoideae | Msin | Misin12G209300 | | | |

| | | | | | | |
|---|---|---|---|---|---|---|
| CYP701A8 | Panicoideae | Msin | Misin18G176400 | | | |
| CYP701A8 | Panicoideae | Msin | Misin19G162400 | | | |
| CYP76L1 | Panicoideae | Msin | Misin19G278000 | | | |
| CYP76L1 | Panicoideae | Phal | Pahal.1G039100 | | | |
| CPS4 | Panicoideae | Phal | Pahal.1G276100 | | | |
| CPS4 | Panicoideae | Phal | Pahal.1G276200 | | | |
| CPS4 | Panicoideae | Phal | Pahal.1G276300 | | | |
| CPS4 | Panicoideae | Phal | Pahal.2G213000 | | | |
| CYP99A2/3 | Panicoideae | Phal | Pahal.2G214200 | | | |
| CYP76L1 | Panicoideae | Phal | Pahal.2G284600 | | | |
| CYP701A8 | Panicoideae | Phal | Pahal.4G105800 | | | |
| CYP701A8 | Panicoideae | Phal | Pahal.4G106200 | | | |
| CYP99A2/3 | Panicoideae | Phal | Pahal.6G156100 | | | |
| KSL4 | Panicoideae | Phal | Pahal.7G288100 | | | |
| MAS1/2 | Panicoideae | Phal | Pahal.8G261800 | | | |
| CYP99A2/3 | Panicoideae | Sbic | Sobic.001G338900 | | | |
| CYP99A2/3 | Panicoideae | Sbic | Sobic.002G090900 | | | |
| CYP76L1 | Panicoideae | Sbic | Sobic.002G220500 | | | |
| CPS4 | Panicoideae | Sbic | Sobic.005G161200 | | | |
| CYP99A2/3 | Panicoideae | Sbic | Sobic.005G217500 | | | |
| CYP99A2/3 | Panicoideae | Sbic | Sobic.006G010200 | | | |
| KSL4 | Panicoideae | Sbic | Sobic.006G211500 | | | |
| CYP701A8 | Panicoideae | Sbic | Sobic.010G172700 | | | |
| CPS4 | Panicoideae | Sita | Seita.1G203600 | | | |
| CPS4 | Panicoideae | Sita | Seita.2G144900 | clustered | Sita_c2 | |
| CYP76L1 | Panicoideae | Sita | Seita.2G145700 | clustered | Sita_c2 | |
| CYP99A2/3 | Panicoideae | Sita | Seita.2G145800 | clustered | Sita_c2 | |
| CYP99A2/3 | Panicoideae | Sita | Seita.2G145900 | clustered | Sita_c2 | |
| CYP76L1 | Panicoideae | Sita | Seita.2G224300 | | | |
| CYP701A8 | Panicoideae | Sita | Seita.4G235400 | | | |
| CYP701A8 | Panicoideae | Sita | Seita.4G235500 | | | |
| CYP701A8 | Panicoideae | Sita | Seita.4G235600 | | | |
| CYP99A2/3 | Panicoideae | Sita | Seita.6G023100 | | | |
| CYP99A2/3 | Panicoideae | Sita | Seita.6G100500 | | | |
| CYP99A2/3 | Panicoideae | Sita | Seita.6G101900 | | | |
| CYP99A2/3 | Panicoideae | Sita | Seita.7G013400 | | | |
| KSL4 | Panicoideae | Sita | Seita.7G233600 | | | |
| CPS4 | Panicoideae | Svir | Sevir.1G206900 | | | |
| CPS4 | Panicoideae | Svir | Sevir.2G150600 | clustered | Svir_c2 | |
| CYP76L1 | Panicoideae | Svir | Sevir.2G151800 | clustered | Svir_c2 | |
| CYP99A2/3 | Panicoideae | Svir | Sevir.2G152000 | clustered | Svir_c2 | |
| CYP99A2/3 | Panicoideae | Svir | Sevir.2G152300 | clustered | Svir_c2 | |
| CYP76L1 | Panicoideae | Svir | Sevir.2G234640 | | | |
| CYP701A8 | Panicoideae | Svir | Sevir.4G247600 | | | |
| CYP701A8 | Panicoideae | Svir | Sevir.4G247700 | | | |
| CYP701A8 | Panicoideae | Svir | Sevir.4G247800 | | | |
| CYP99A2/3 | Panicoideae | Svir | Sevir.6G021400 | | | |
| CYP99A2/3 | Panicoideae | Svir | Sevir.6G110300 | | | |
| CYP99A2/3 | Panicoideae | Svir | Sevir.6G112300 | | | |
| CYP99A2/3 | Panicoideae | Svir | Sevir.7G042500 | | | |
| KSL4 | Panicoideae | Svir | Sevir.7G245200 | | | |
| CYP99A2/3 | Panicoideae | Svir | Sevir.9G273001 | | | |
| CYP99A2/3 | Panicoideae | Zmay | Zm00008a001750 | | | |
| KSL4 | Panicoideae | Zmay | Zm00008a006308 | | | |
| CYP99A2/3 | Panicoideae | Zmay | Zm00008a015052 | clustered | | |
| CPS4 | Panicoideae | Zmay | Zm00008a015054 | clustered | | |
| CPS4 | Panicoideae | Zmay | Zm00008a015060 | clustered | | |
| CYP99A2/3 | Panicoideae | Zmay | Zm00008a015061 | clustered | | |
| CYP701A8 | Panicoideae | Zmay | Zm00008a020723 | | | |
| CYP76L1 | Panicoideae | Zmay | Zm00008a028208 | | | |
| CYP701A8 | Panicoideae | Zmay | Zm00008a034463 | | | |
| CYP701A8 | Panicoideae | Zmay | Zm00008a034464 | | | |
| CYP99A2/3 | Panicoideae | Zmay | Zm00008a035392 | | | |
| CPS4 | Panicoideae | Zmay | Zm00008a037235 | paired | | |
| KSL4 | Panicoideae | Zmay | Zm00008a037236 | paired | | |
| CYP99A2/3 | Panicoideae | Zmay | Zm00008a037518 | | | |
| KSL4 | Oryzoideae | Lper | LPERR02G17300 | paired | Lper_c2BGC | c2BGC |
| CPS4 | Oryzoideae | Lper | LPERR02G17320 | paired | Lper_c2BGC | c2BGC |
| KSL4 | Oryzoideae | Lper | LPERR04G20550 | | | |
| CYP701A8 | Oryzoideae | Lper | LPERR06G15600 | | | |
| CYP701A8 | Oryzoideae | Lper | LPERR06G15610 | | | |
| CYP701A8 | Oryzoideae | Lper | LPERR06G15620 | | | |
| CYP76L1 | Oryzoideae | Lper | LPERR09G08730 | | | |
| KSL4 | Oryzoideae | Obar | OBART02G21250.1 | clustered | Obar_c2BGC | Phytoalexins c2BGC |
| CPS4 | Oryzoideae | Obar | OBART02G21280.1 | clustered | Obar_c2BGC | Phytoalexins c2BGC |
| CPS4 | Oryzoideae | Obar | OBART04G02860.1 | clustered | Obar_MABGC | MABGC |
| CYP99A2/3 | Oryzoideae | Obar | OBART04G02870.1 | clustered | Obar_MABGC | MABGC |

| | | | | | | | |
|---|---|---|---|---|---|---|---|
| MAS1/2 | Oryzoideae | Obar | OBART04G02910.1 | clustered | Obar_MABGC | | MABGC |
| KSL4 | Oryzoideae | Obar | OBART04G02920.1 | clustered | Obar_MABGC | | MABGC |
| CYP99A2/3 | Oryzoideae | Obar | OBART04G02940.1 | clustered | Obar_MABGC | | MABGC |
| KSL4 | Oryzoideae | Obar | OBART04G25420.1 | | | | |
| CYP701A8 | Oryzoideae | Obar | OBART06G18930.1 | TD | | | |
| CYP701A8 | Oryzoideae | Obar | OBART06G18950.1 | TD | | | |
| CYP701A8 | Oryzoideae | Obar | OBART06G18970.1 | TD | | | |
| CYP701A8 | Oryzoideae | Obar | OBART06G18990.1 | TD | | | |
| CYP701A8 | Oryzoideae | Obar | OBART06G19000.1 | TD | | | |
| CPS4 | Oryzoideae | Obar | OBART09G05090.1 | | | | |
| CPS4 | Oryzoideae | Obar | OBART09G05170.1 | | | | |
| CYP76L1 | Oryzoideae | Obar | OBART09G12160.1 | | | | |
| CYP99A2/3 | Oryzoideae | Obra | OB0266G10010.1 | | | | |
| CYP99A2/3 | Oryzoideae | Obra | OB04G12330.1 | | | | |
| CYP99A2/3 | Oryzoideae | Obra | OB04G12340.1 | | | | |
| KSL4 | Oryzoideae | Obra | OB04G31870.1 | | | | |
| CYP99A2/3 | Oryzoideae | Obra | OB06G23390.1 | | | | |
| CYP701A8 | Oryzoideae | Obra | OB06G26530.1 | TD | | | |
| CYP701A8 | Oryzoideae | Obra | OB06G26540.1 | TD | | | |
| CYP701A8 | Oryzoideae | Obra | OB06G26550.1 | TD | | | |
| CYP701A8 | Oryzoideae | Obra | OB06G26560.1 | TD | | | |
| CYP701A8 | Oryzoideae | Obra | OB06G26570.1 | TD | | | |
| CYP76L1 | Oryzoideae | Obra | OB09G18690.1 | | | | |
| CPS4 | Oryzoideae | Obra | OB11G20480.1 | | | | |
| KSL4 | Oryzoideae | Ogla | ORGLA02G0183000.1 | clustered | Ogla_c2BGC | | Phytoalexins c2BGC |
| CPS4 | Oryzoideae | Ogla | ORGLA02G0183400.1 | clustered | Ogla_c2BGC | | Phytoalexins c2BGC |
| CPS4 | Oryzoideae | Ogla | ORGLA04G0020300.1 | clustered | Ogla_MABGC | | MABGC |
| CYP99A2/3 | Oryzoideae | Ogla | ORGLA04G0020400.1 | clustered | Ogla_MABGC | | MABGC |
| MAS1/2 | Oryzoideae | Ogla | ORGLA04G0020700.1 | clustered | Ogla_MABGC | | MABGC |
| MAS1/2 | Oryzoideae | Ogla | ORGLA04G0020800.1 | clustered | Ogla_MABGC | | MABGC |
| KSL4 | Oryzoideae | Ogla | ORGLA04G0021100.1 | clustered | Ogla_MABGC | | MABGC |
| CYP99A2/3 | Oryzoideae | Ogla | ORGLA04G0021400.1 | clustered | Ogla_MABGC | | MABGC |
| KSL4 | Oryzoideae | Ogla | ORGLA04G0220500.1 | | | | |
| CYP701A8 | Oryzoideae | Ogla | ORGLA06G0157100.1 | TD | | | |
| CYP701A8 | Oryzoideae | Ogla | ORGLA06G0157300.1 | TD | | | |
| CYP701A8 | Oryzoideae | Ogla | ORGLA06G0157400.1 | TD | | | |
| CYP701A8 | Oryzoideae | Ogla | ORGLA06G0157500.1 | TD | | | |
| CPS4 | Oryzoideae | Ogla | ORGLA09G0038000.1 | | | | |
| KSL4 | Oryzoideae | Opun | OPUNC02G19210.1 | | | | |
| CPS4 | Oryzoideae | Opun | OPUNC04G02770.1 | clustered | Opun_MABGC | | MABGC |
| CYP99A2/3 | Oryzoideae | Opun | OPUNC04G02780.1 | clustered | Opun_MABGC | | MABGC |
| MAS1/2 | Oryzoideae | Opun | OPUNC04G02800.1 | clustered | Opun_MABGC | | MABGC |
| MAS1/2 | Oryzoideae | Opun | OPUNC04G02820.1 | clustered | Opun_MABGC | | MABGC |
| KSL4 | Oryzoideae | Opun | OPUNC04G02860.1 | clustered | Opun_MABGC | | MABGC |
| CYP99A2/3 | Oryzoideae | Opun | OPUNC04G02870.1 | clustered | Opun_MABGC | | MABGC |
| KSL4 | Oryzoideae | Opun | OPUNC04G22480.1 | | | | |
| CYP701A8 | Oryzoideae | Opun | OPUNC06G17160.1 | TD | | | |
| CYP701A8 | Oryzoideae | Opun | OPUNC06G17180.1 | TD | | | |
| CYP76L1 | Oryzoideae | Opun | OPUNC09G10670.1 | | | | |
| CPS4 | Oryzoideae | Opun | OPUNC10G06550.1 | | | | |
| KSL4 | Oryzoideae | Oruf | ORUFI02G22350.1 | clustered | Oruf_c2BGC | | Phytoalexins c2BGC |
| CPS4 | Oryzoideae | Oruf | ORUFI02G22400.1 | clustered | Oruf_c2BGC | | Phytoalexins c2BGC |
| CPS4 | Oryzoideae | Oruf | ORUFI04G03540.1 | clustered | Oruf_MABGC | | MABGC |
| CYP99A2/3 | Oryzoideae | Oruf | ORUFI04G03550.1 | clustered | Oruf_MABGC | | MABGC |
| MAS1/2 | Oryzoideae | Oruf | ORUFI04G03580.1 | clustered | Oruf_MABGC | | MABGC |
| MAS1/2 | Oryzoideae | Oruf | ORUFI04G03590.1 | clustered | Oruf_MABGC | | MABGC |
| KSL4 | Oryzoideae | Oruf | ORUFI04G03600.1 | clustered | Oruf_MABGC | | MABGC |
| CYP99A2/3 | Oryzoideae | Oruf | ORUFI04G03620.1 | clustered | Oruf_MABGC | | MABGC |
| KSL4 | Oryzoideae | Oruf | ORUFI04G26630.1 | clustered | Oruf_MABGC | | MABGC |
| CYP701A8 | Oryzoideae | Oruf | ORUFI06G20020.1 | TD | | | |
| CYP701A8 | Oryzoideae | Oruf | ORUFI06G20040.1 | TD | | | |
| CYP701A8 | Oryzoideae | Oruf | ORUFI06G20070.1 | TD | | | |
| CYP701A8 | Oryzoideae | Oruf | ORUFI06G20090.1 | TD | | | |
| CYP701A8 | Oryzoideae | Oruf | ORUFI06G20100.1 | TD | | | |
| CPS4 | Oryzoideae | Oruf | ORUFI09G05890.1 | | | | |
| CYP76L1 | Oryzoideae | Oruf | ORUFI09G13060.1 | | | | |
| KSL4 | Oryzoideae | Osat | LOC_Os02g36140.2 | clustered | Osat_c2 | KSL7 | Phytoalexins c2BGC |
| CPS4 | Oryzoideae | Osat | LOC_Os02g36210 | clustered | Osat_c2 | CPS2 | Phytoalexins c2BGC |
| CPS4 | Oryzoideae | Osat | LOC_Os04g09900 | clustered | Osat_MABGC | | MABGC |
| CYP99A2/3 | Oryzoideae | Osat | LOC_Os04g09920 | clustered | Osat_MABGC | | MABGC |
| MAS1/2 | Oryzoideae | Osat | LOC_Os04g10000 | clustered | Osat_MABGC | | MABGC |
| MAS1/2 | Oryzoideae | Osat | LOC_Os04g10010 | clustered | Osat_MABGC | | MABGC |
| KSL4 | Oryzoideae | Osat | LOC_Os04g10060 | clustered | Osat_MABGC | | MABGC |
| CYP99A2/3 | Oryzoideae | Osat | LOC_Os04g10160 | clustered | Osat_MABGC | | MABGC |
| KSL4 | Oryzoideae | Osat | LOC_Os04g52210 | | | KSL3 | |
| KSL4 | Oryzoideae | Osat | LOC_Os04g52230 | | | KS1 | |
| CYP701A8 | Oryzoideae | Osat | LOC_Os06g37224 | TD | | | |

| | | | | | | |
|---|---|---|---|---|---|---|
| CYP701A8 | Oryzoideae | Osat | LOC_Os06g37300 | TD | | |
| CYP701A8 | Oryzoideae | Osat | LOC_Os06g37330 | TD | | |
| CYP701A8 | Oryzoideae | Osat | LOC_Os06g37364 | TD | | |
| CPS4 | Oryzoideae | Osat | LOC_Os09g15050 | | | |
| CYP76L1 | Oryzoideae | Osat | LOC_Os09g27500 | | | |
| CYP701A8 | Oryzoideae | Zlat | 10003144 | | | |
| CYP76L1 | Oryzoideae | Zlat | 10012738 | | | |
| KSL4 | Oryzoideae | Zlat | 10013824 | clustered | Zlat_c2BGC | Phytoalexins c2BGC |
| CPS4 | Oryzoideae | Zlat | 10013826 | clustered | Zlat_c2BGC | Phytoalexins c2BGC |
| CPS4 | Oryzoideae | Zlat | 10013827 | clustered | Zlat_c2BGC | Phytoalexins c2BGC |
| CPS4 | Oryzoideae | Zlat | 10013829 | clustered | Zlat_c2BGC | Phytoalexins c2BGC |
| CYP701A8 | Oryzoideae | Zlat | 10014669 | TD | | |
| CYP701A8 | Oryzoideae | Zlat | 10014671 | TD | | |
| CYP701A8 | Oryzoideae | Zlat | 10014672 | TD | | |
| CYP701A8 | Oryzoideae | Zlat | 10036611 | | | |
| KSL4 | Oryzoideae | Zlat | 10041552 | | | |
| CPS4 | Chloridoideae | Cson | CsA102785.1 | | | |
| CYP76L1 | Chloridoideae | Cson | CsA302295.1 | | | |
| CYP701A8 | Chloridoideae | Cson | CsA302831.1 | | | |
| KSL4 | Chloridoideae | Cson | CsA500576.1 | | | |
| CYP99A2/3 | Chloridoideae | Cson | CsA502071.1 | | | |
| CYP99A2/3 | Chloridoideae | Cson | CsA600455.1 | paired | | |
| KSL4 | Chloridoideae | Cson | CsA600456.1 | paired | | |
| MAS1/2 | Chloridoideae | Cson | CsA600829.1 | | | |
| CYP99A2/3 | Chloridoideae | Cson | CsB101450.1 | | | |
| KSL4 | Chloridoideae | Cson | CsB200568.1 | | | |
| CYP76L1 | Chloridoideae | Cson | CsB302343.1 | | | |
| CYP701A8 | Chloridoideae | Cson | CsB302888.1 | | | |
| CPS4 | Chloridoideae | Cson | CsB601177.1 | paired | | |
| MAS1/2 | Chloridoideae | Cson | CsB601178.1 | paired | | |
| KSL4 | Chloridoideae | Ecur | TVT97615 | | | |
| CPS4 | Chloridoideae | Ecur | TVT99217 | | | |
| CYP99A2/3 | Chloridoideae | Ecur | TVU01138 | | | |
| CYP701A8 | Chloridoideae | Ecur | TVU07937 | | | |
| CYP701A8 | Chloridoideae | Ecur | TVU07938 | | | |
| CYP76L1 | Chloridoideae | Ecur | TVU09766 | | | |
| KSL4 | Chloridoideae | Ecur | TVU15791 | | | |
| KSL4 | Chloridoideae | Ecur | TVU15792 | | | |
| KSL4 | Chloridoideae | Ecur | TVU15793 | | | |
| CPS4 | Chloridoideae | Ecur | TVU29579 | | | |
| CPS4 | Chloridoideae | Ecur | TVU29590 | | | |
| CPS4 | Chloridoideae | Ecur | TVU29591 | | | |
| CPS4 | Chloridoideae | Ecur | TVU29592 | | | |
| CPS4 | Chloridoideae | Ecur | TVU29601 | | | |
| CPS4 | Chloridoideae | Ecur | TVU30191 | | | |
| CYP99A2/3 | Chloridoideae | Ecur | TVU30765 | | | |
| CYP99A2/3 | Chloridoideae | Ecur | TVU30766 | | | |
| CYP99A2/3 | Chloridoideae | Ecur | TVU30767 | | | |
| CPS4 | Chloridoideae | Ecur | TVU33980 | | | |
| CPS4 | Chloridoideae | Ecur | TVU33986 | | | |
| CPS4 | Chloridoideae | Ecur | TVU42376 | | | |
| CPS4 | Chloridoideae | Ecur | TVU42379 | | | |
| CYP99A2/3 | Chloridoideae | Ecur | TVU42411 | clustered | Ecur_MABGC | MABGC |
| MAS1/2 | Chloridoideae | Ecur | TVU42414 | clustered | Ecur_MABGC | MABGC |
| CPS4 | Chloridoideae | Ecur | TVU42419 | clustered | Ecur_MABGC | MABGC |
| CPS4 | Chloridoideae | Ecur | TVU42420 | clustered | Ecur_MABGC | MABGC |
| CYP99A2/3 | Chloridoideae | Ecur | TVU42421 | clustered | Ecur_MABGC | MABGC |
| CYP99A2/3 | Chloridoideae | Ecur | TVU42422 | clustered | Ecur_MABGC | MABGC |
| CYP99A2/3 | Chloridoideae | Ecur | TVU42424 | clustered | Ecur_MABGC | MABGC |
| KSL4 | Chloridoideae | Ecur | TVU42425 | clustered | Ecur_MABGC | MABGC |
| CPS4 | Chloridoideae | Ecur | TVU42426 | clustered | Ecur_MABGC | MABGC |
| CYP99A2/3 | Chloridoideae | Ecur | TVU51751 | | | |
| CYP99A2/3 | Chloridoideae | Ecur | TVU51752 | | | |
| CPS4 | Chloridoideae | Enin | En_0020078-RA | clustered | Enin_c002 | |
| CYP99A2/3 | Chloridoideae | Enin | En_0020079-RA | clustered | Enin_c002 | |
| CYP99A2/3 | Chloridoideae | Enin | En_0020081-RA | clustered | Enin_c002 | |
| CPS4 | Chloridoideae | Enin | En_0020085-RA | clustered | Enin_c002 | |
| CYP99A2/3 | Chloridoideae | Enin | En_0020092-RA | clustered | Enin_c002 | |
| MAS1/2 | Chloridoideae | Enin | En_0020097-RA | clustered | Enin_c002 | |
| CYP76L1 | Chloridoideae | Enin | En_0029726-RA | | | |
| CYP99A2/3 | Chloridoideae | Enin | En_0036858-RA | | | |
| CPS4 | Chloridoideae | Enin | En_0054442-RA | | | |
| CPS4 | Chloridoideae | Enin | En_0055080-RA | | | |
| CPS4 | Chloridoideae | Enin | En_0055081-RA | | | |
| CYP76L1 | Chloridoideae | Enin | En_0057220-RA | | | |
| CYP76L1 | Chloridoideae | Enin | En_0063575-RA | | | |
| MAS1/2 | Chloridoideae | Enin | En_0064379-RA | | | |

| | | | | |
|---|---|---|---|---|
| KSL4 | Chloridoideae | Enin | En_0066312-RA | |
| KSL4 | Chloridoideae | Enin | En_0066313-RA | |
| KSL4 | Chloridoideae | Enin | En_0091563-RA | |
| CYP99A2/3 | Chloridoideae | Enin | En_0095132-RA | |
| CYP99A2/3 | Chloridoideae | Enin | En_0102186-RA | paired |
| CPS4 | Chloridoideae | Enin | En_0102188-RA | paired |
| CYP99A2/3 | Chloridoideae | Enin | En_0105784-RA | paired |
| CPS4 | Chloridoideae | Enin | En_0105786-RA | paired |
| CYP99A2/3 | Chloridoideae | Enin | En_0109695-RA | |
| CYP99A2/3 | Chloridoideae | Enin | En_0109696-RA | |
| CYP99A2/3 | Chloridoideae | Enin | En_0109700-RA | |
| CYP99A2/3 | Chloridoideae | Etef | Et_10000679 | |
| CYP99A2/3 | Chloridoideae | Etef | Et_10000682 | |
| CPS4 | Chloridoideae | Etef | Et_10000876 | |
| CYP99A2/3 | Chloridoideae | Etef | Et_10002828 | |
| CYP99A2/3 | Chloridoideae | Etef | Et_10002831 | |
| CPS4 | Chloridoideae | Etef | Et_1005482 | |
| CYP99A2/3 | Chloridoideae | Etef | Et_1006263 | |
| CYP99A2/3 | Chloridoideae | Etef | Et_1006300 | |
| CYP99A2/3 | Chloridoideae | Etef | Et_1006301 | |
| CYP99A2/3 | Chloridoideae | Etef | Et_1011181 | |
| CYP99A2/3 | Chloridoideae | Etef | Et_1011182 | |
| CYP99A2/3 | Chloridoideae | Etef | Et_1011183 | |
| CPS4 | Chloridoideae | Etef | Et_1013596 | |
| CYP76L1 | Chloridoideae | Etef | Et_2016179 | |
| CYP76L1 | Chloridoideae | Etef | Et_2020290 | |
| KSL4 | Chloridoideae | Etef | Et_7052091 | |
| KSL4 | Chloridoideae | Etef | Et_7054730 | |
| CYP99A2/3 | Chloridoideae | Etef | Et_8056081 | |
| CPS4 | Chloridoideae | Etef | Et_8056235 | |
| MAS1/2 | Chloridoideae | Etef | Et_8056826 | paired |
| CYP99A2/3 | Chloridoideae | Etef | Et_8056829 | paired |
| CPS4 | Chloridoideae | Etef | Et_8058067 | |
| MAS1/2 | Chloridoideae | Etef | Et_8058544 | |
| CPS4 | Chloridoideae | Etef | Et_8059177 | |
| CYP99A2/3 | Chloridoideae | Etef | Et_8059196 | |
| CYP99A2/3 | Chloridoideae | Etef | Et_8059202 | |
| CYP99A2/3 | Chloridoideae | Etef | Et_8060427 | paired |
| MAS1/2 | Chloridoideae | Etef | Et_8060433 | paired |
| CPS4 | Chloridoideae | Otho | Oropetium 20150105 02521 | |
| CYP701A8 | Chloridoideae | Otho | Oropetium_20150105_17592 | |
| CYP701A8 | Chloridoideae | Zjap | Zjn_sc00008.1.g06820.1.sm.mk | |
| CPS4 | Chloridoideae | Zjap | Zjn_sc00017.1.g02660.1.am.mk | |
| CPS4 | Chloridoideae | Zjap | Zjn_sc00020.1.g00450.1.sm.mk | |
| CPS4 | Chloridoideae | Zjap | Zjn_sc00020.1.g00460.1.sm.mk | |
| CYP99A2/3 | Chloridoideae | Zjap | Zjn_sc00033.1.g00200.1.am.mk | |
| CYP76L1 | Chloridoideae | Zjap | Zjn_sc00039.1.g01660.1.sm.mk | |
| CPS4 | Bambusoideae | Bamp | Bam004236.1 | |
| CYP76L1 | Bambusoideae | Bamp | Bam027633.1 | |
| KSL4 | Bambusoideae | Bamp | Bam034720.1 | |
| CYP99A2/3 | Bambusoideae | Bamp | Bam045979.1 | |
| CYP701A8 | Bambusoideae | Olat | Ola003381.1 | TD |
| CYP701A8 | Bambusoideae | Olat | Ola003387.1 | TD |
| CYP701A8 | Bambusoideae | Olat | Ola003391.1 | TD |
| CYP701A8 | Bambusoideae | Olat | Ola003395.1 | TD |
| CPS4 | Bambusoideae | Olat | Ola021372.1 | |
| CPS4 | Bambusoideae | Olat | Ola021374.1 | |
| CPS4 | Bambusoideae | Olat | Ola021375.1 | |
| CPS4 | Bambusoideae | Olat | Ola021376.1 | |
| KSL4 | Bambusoideae | Olat | Ola022358.1 | |
| KSL4 | Bambusoideae | Olat | Ola022368.2 | |
| CYP76L1 | Bambusoideae | Olat | Ola025389.1 | |

Table S3. Transcriptomic profiling of wheat under infection of pathogens

| Cluster | Cluster c2_2 | | | | | | | | | | | | | | | | | |
|---|---|---|---|---|---|---|---|---|---|---|---|---|---|---|---|---|---|---|
| Subgenome | A | | | | B | | | | | | | | | | D | | | |
| Gene Locus | TraesCS2A02G42450 | TraesCS2A02G42510 | TraesCS2A02G42520 | TraesCS2A02G42540 | TraesCS2B02G44410 | TraesCS2B02G44490 | TraesCS2B02G44500 | TraesCS2B02G44510 | TraesCS2B02G44520 | TraesCS2B01G44530 | TraesCS2B01G44540 | TraesCS2B02G44550 | TraesCS2B02G44560 | TraesCS2B02G44570 | TraesCS2D02G42230 | TraesCS2D02G42310 | TraesCS2D02G42320 | TraesCS2D02G42330 |
| Gene Type | CYP99 | KSL | CYP99 | KSL | CYP99 | KSL | CYP99 | KSL | KSL | CYP70 | CYP70 | CPS | CYP99 | KSL | CYP99 | KSL | CYP99 | KSL |
| **Data FHB 2DL (doi: 10.3389/fpls.2018.00037). Tissue: Spikelet (SP) and Rachis (RACH); Material: 2618, 2890; Treatment: control (H2O), *Fusarium graminearum* inoculation (FG)** | | | | | | | | | | | | | | | | | | |
| 2618_H2O_RACH | 0.00 | 0.00 | 0.12 | 3.46 | 0.00 | 0.02 | 0.00 | 0.53 | 0.12 | 0.80 | 0.10 | 0.31 | 0.61 | 1.54 | 0.00 | 0.00 | 0.82 | 2.39 |
| 2618_FG_RACH | 0.02 | 0.00 | 0.08 | 0.63 | 0.03 | 0.50 | 0.00 | 23.91 | 14.81 | 75.60 | 1.60 | ##### | ##### | 0.46 | 0.01 | 0.01 | 0.10 | 0.63 |
| 2618_H2O_SP | 0.01 | 0.00 | 1.83 | 1.62 | 0.00 | 0.01 | 0.02 | 2.07 | 0.83 | 3.70 | 0.20 | 1.66 | 2.69 | 0.81 | 0.00 | 0.01 | 0.06 | 0.93 |
| 2618_FG_SP | 0.03 | 0.01 | 0.86 | 1.17 | 0.00 | 0.29 | 0.00 | 15.68 | 12.63 | 66.20 | 3.00 | ##### | ##### | 0.59 | 0.02 | 0.00 | 0.01 | 0.64 |
| 2890_H2O_SP | 0.01 | 0.00 | 1.64 | 1.73 | 0.00 | 0.00 | 0.00 | 0.50 | 0.17 | 0.80 | 0.20 | 0.38 | 0.64 | 1.03 | 0.00 | 0.00 | 0.06 | 1.68 |
| 2890_FG_SP | 0.04 | 0.00 | 0.49 | 0.43 | 0.11 | 0.23 | 0.00 | 5.41 | 10.38 | 86.00 | 9.50 | ##### | ##### | 0.21 | 0.02 | 0.00 | 0.01 | 0.49 |
| 2890_H2O_RACH | 0.00 | 0.00 | 0.11 | 3.25 | 0.01 | 0.01 | 0.00 | 2.38 | 0.85 | 3.20 | 0.10 | 2.54 | 3.58 | 1.34 | 0.00 | 0.00 | 0.47 | 3.22 |
| 2890_FG_RACH | 0.15 | 0.00 | 0.01 | 0.10 | 0.00 | 0.11 | 0.00 | 4.98 | 5.58 | 53.40 | 4.40 | 84.77 | ##### | 0.09 | 0.02 | 0.00 | 0.00 | 0.28 |
| **Data RNAseq on *Fusarium pseudograminearum* infected wheat (doi: 10.1111/pbi.12651). Tissue: coleoptile sheath; Material: Chara; Treatment: control (Mock), *Fusarium pseudograminearum* infection (Fp)** | | | | | | | | | | | | | | | | | | |
| Chara_Mock | 0.00 | 0.00 | 5.75 | 2.91 | 0.01 | 0.00 | 0.00 | 0.37 | 0.08 | 0.50 | 0.10 | 0.11 | 0.23 | 1.95 | 0.13 | 0.00 | 0.13 | 4.59 |
| Chara_Fp | 0.00 | 0.00 | 5.43 | 2.40 | 0.00 | 0.00 | 0.00 | 0.90 | 1.77 | 6.00 | 0.20 | 2.59 | 5.84 | 1.73 | 0.18 | 0.00 | 0.13 | 5.28 |
| **Data Powdery Mildew Pathogen Stress (doi: 10.1186/1471-2164-15-898). Tissue: Leaf. Material: N9134; Treatment: powdery mildew (*Blumeria graminis*) infection.** | | | | | | | | | | | | | | | | | | |
| non-innoculation | 0.03 | 0.00 | 0.70 | 0.30 | 0.00 | 0.00 | 0.00 | 0.32 | 0.03 | 1.20 | 0.20 | 0.18 | 0.59 | 0.69 | 0.07 | 0.00 | 0.18 | 1.94 |
| Powdery_24h | 0.00 | 0.00 | 0.74 | 0.26 | 0.01 | 0.00 | 0.00 | 80.81 | 1.36 | ##### | 15.60 | 30.17 | 99.25 | 0.51 | 0.00 | 0.00 | 0.05 | 1.75 |
| Powdery_48h | 0.00 | 0.04 | 1.02 | 0.69 | 0.00 | 0.00 | 0.00 | 28.84 | 0.26 | ##### | 9.30 | 16.62 | 36.37 | 1.08 | 0.14 | 0.00 | 0.05 | 3.11 |
| Powdery_72h | 0.03 | 0.00 | 0.70 | 0.83 | 0.00 | 0.00 | 0.00 | 3.01 | 0.03 | 11.20 | 1.00 | 2.08 | 4.25 | 1.10 | 0.07 | 0.00 | 0.10 | 2.92 |
| **Data *Pyrenophora tritici-repentis* inoculation. Tissue: Leaf. Material: Glenlea, Salamouni; Treatment: *Pyrenophora tritici-repentis* inoculation** | | | | | | | | | | | | | | | | | | |
| Glenlea_Control_48pi | 0.00 | 0.00 | 0.19 | 1.09 | 0.00 | 0.00 | 0.00 | 0.35 | 0.00 | 0.30 | 0.10 | 0.00 | 0.05 | 0.94 | 0.00 | 0.00 | 0.30 | 3.40 |
| Glenlea_Fungus_48pi | 0.00 | 0.00 | 0.25 | 0.42 | 0.04 | 0.04 | 0.00 | 14.09 | 19.13 | 80.10 | 5.20 | 59.47 | ##### | 1.34 | 0.00 | 0.00 | 0.31 | 2.65 |
| Salamouni_Control_48pi | 0.00 | 0.00 | 0.09 | 0.72 | 0.00 | 0.00 | 0.00 | 0.42 | 0.43 | 1.10 | 0.00 | 0.23 | 0.09 | 1.31 | 0.00 | 0.00 | 0.52 | 2.52 |
| Salamouni_Fungus_48pi | 0.00 | 0.00 | 0.11 | 0.68 | 0.00 | 0.08 | 0.00 | 87.31 | 91.63 | ##### | 1.20 | ##### | ##### | 1.16 | 0.00 | 0.00 | 0.08 | 1.98 |
| **Data Stripe rust pathogen stress (doi: 10.1186/1471-2164-15-898). Tissue: Leaf. Material: N9134. Treatment: Stripe rust (*Puccinia striiformis*) infection.** | | | | | | | | | | | | | | | | | | |
| non-innoculation | 0.03 | 0.00 | 0.70 | 0.30 | 0.00 | 0.00 | 0.00 | 0.32 | 0.03 | 1.20 | 0.20 | 0.18 | 0.59 | 0.69 | 0.07 | 0.00 | 0.18 | 1.94 |
| Stripe24h | 0.03 | 0.00 | 1.56 | 0.36 | 0.00 | 0.00 | 0.00 | 0.59 | 0.00 | 4.30 | 0.50 | 0.22 | 1.22 | 0.46 | 0.06 | 0.00 | 0.12 | 2.47 |
| Stripe48h | 0.00 | 0.00 | 1.27 | 0.43 | 0.00 | 0.00 | 0.00 | 0.48 | 0.01 | 2.20 | 0.20 | 0.13 | 0.46 | 0.81 | 0.14 | 0.00 | 0.23 | 2.12 |
| Stripe72h | 0.00 | 0.00 | 1.00 | 0.28 | 0.00 | 0.00 | 0.00 | 0.36 | 0.03 | 2.00 | 0.10 | 0.30 | 0.88 | 0.54 | 0.10 | 0.00 | 0.31 | 3.07 |
| **Data *Xanthomonas translucens* infection. Tissue: Root, Leaf. Material: Chinese Spring. Treatment: *Xanthomonas translucens* infection.** | | | | | | | | | | | | | | | | | | |
| Control_Root | 0.01 | 0.64 | 2.12 | 9.24 | 0.00 | 0.02 | 2.08 | 0.18 | 0.14 | 0.10 | 0.00 | 0.39 | 0.16 | 14.98 | 0.01 | 0.04 | 18.27 | 10.29 |
| Xt_Root | 0.03 | 1.00 | 1.82 | 9.34 | 0.01 | 0.00 | 1.92 | 0.41 | 0.08 | 0.70 | 0.10 | 0.59 | 1.06 | 15.14 | 0.06 | 0.13 | 14.15 | 9.63 |
| Control_Leaf | 0.00 | 0.00 | 0.01 | 0.83 | 0.00 | 0.00 | 0.00 | 0.05 | 0.00 | 0.10 | 0.00 | 0.02 | 0.00 | 0.55 | 0.00 | 0.00 | 0.02 | 0.71 |
| Xt_Leaf | 0.00 | 0.00 | 0.02 | 0.38 | 0.01 | 0.00 | 0.00 | 0.30 | 0.08 | 2.20 | 0.00 | 0.91 | 1.70 | 0.30 | 0.00 | 0.00 | 0.03 | 0.49 |

| Cluster | Cluster c2_1 | | | | | | | | | | | | | | | | | | | | | | |
|---|---|---|---|---|---|---|---|---|---|---|---|---|---|---|---|---|---|---|---|---|---|---|---|
| Subgenome | A | | | | | | | | B | | | | | D | | | | | | | | | |
| Gene Locus | TraesCS2A02G02710 | TraesCS2A02G02720 | TraesCS2A02G02740 | TraesCS2A02G02760 | TraesCS2A02G02770 | TraesCS2A02G02780 | TraesCS2A02G02810 | TraesCS2A02G02820 | TraesCS2B02G04040 | TraesCS2B02G04080 | TraesCS2B02G04090 | TraesCS2B02G04100 | TraesCS2B02G04110 | TraesCS2D02G02940 | TraesCS2D02G02950 | TraesCS2D02G02960 | TraesCS2D02G02970 | TraesCS2D02G02990 | TraesCS2D02G03000 | TraesCS2D02G03010 | TraesCS2D02G03020 | TraesCS2D02G03030 | TraesCS2D02G03070 |
| Gene Type | CPS | CPS | CYP99 | KSL | CYP99 | CPS | KSL | CPS | CPS | CPS | KSL | KSL | CPS | KSL | CYP99 | CPS | CYP99 | CYP99 | CYP99 | KSL | KSL | CPS | CYP99 |
| 2618_H2O_RACH | 0.00 | 0.01 | 0.03 | 0.02 | 0.09 | 0.23 | 0.00 | 0.00 | 0.04 | 0.00 | 0.00 | 0.00 | 0.00 | 0.04 | 0.09 | 0.09 | 0.29 | 0.12 | 0.03 | 0.09 | 0.02 | 0.00 | 0.00 |
| 2618_FG_RACH | 0.00 | 2.13 | 30.81 | 32.48 | ##### | 26.73 | 0.00 | 0.00 | 0.96 | 0.00 | 0.00 | 0.00 | 0.00 | 8.37 | 72.80 | 28.78 | 40.79 | 20.14 | 11.70 | 9.49 | 0.00 | 0.01 | 0.00 |
| 2618_H2O_SP | 0.74 | 0.01 | 0.03 | 0.01 | 0.06 | 0.73 | 0.02 | 0.00 | 0.45 | 0.00 | 0.00 | 0.00 | 0.00 | 0.01 | 0.02 | 0.34 | 0.90 | 0.66 | 0.39 | 0.28 | 0.00 | 0.00 | 0.00 |
| 2618_FG_SP | 0.21 | 0.09 | 4.86 | 6.20 | 10.96 | 29.47 | 0.00 | 0.00 | 1.71 | 0.00 | 0.00 | 0.00 | 0.00 | 1.14 | 12.24 | 33.59 | 18.59 | 5.93 | 3.07 | 5.60 | 0.00 | 0.00 | 0.00 |
| 2890_H2O_SP | 0.42 | 0.00 | 0.00 | 0.02 | 0.02 | 0.16 | 0.01 | 0.00 | 0.03 | 0.00 | 0.00 | 0.00 | 0.00 | 0.00 | 0.04 | 0.13 | 0.37 | 0.12 | 0.12 | 0.08 | 0.01 | 0.00 | 0.00 |
| 2890_FG_SP | 0.02 | 0.03 | 6.55 | 5.97 | 8.97 | 34.06 | 0.00 | 0.00 | 0.12 | 0.00 | 0.00 | 0.00 | 0.00 | 0.96 | 16.64 | 47.40 | 13.89 | 3.99 | 1.41 | 3.36 | 0.01 | 0.01 | 0.01 |
| 2890_H2O_RACH | 0.00 | 0.00 | 0.11 | 0.10 | 0.58 | 0.86 | 0.00 | 0.00 | 0.37 | 0.00 | 0.00 | 0.00 | 0.00 | 0.03 | 0.33 | 0.43 | 0.91 | 0.81 | 0.34 | 0.30 | 0.03 | 0.00 | 0.00 |
| 2890_FG_RACH | 0.00 | 1.54 | 25.86 | 22.99 | 91.79 | 12.10 | 0.00 | 0.00 | 0.31 | 0.02 | 0.00 | 0.00 | 0.00 | 6.67 | 61.64 | 17.86 | 19.10 | 4.87 | 2.25 | 2.99 | 0.00 | 0.00 | 0.02 |
| Chara_Mock | 0.00 | 0.00 | 0.00 | 0.01 | 0.10 | 0.03 | 0.00 | 0.00 | 0.00 | 0.00 | 0.00 | 0.00 | 0.00 | 0.00 | 0.02 | 0.02 | 0.03 | 0.02 | 0.02 | 0.06 | 0.04 | 0.00 | 0.00 |
| Chara_Fp | 0.00 | 0.00 | 0.35 | 0.27 | 3.19 | 1.07 | 0.00 | 0.00 | 0.00 | 0.00 | 0.00 | 0.00 | 0.00 | 0.12 | 0.92 | 0.56 | 1.17 | 1.15 | 1.30 | 0.73 | 0.05 | 0.00 | 0.00 |
| non-innoculation | 0.00 | 0.00 | 0.00 | 0.03 | 0.09 | 0.10 | 0.00 | 0.00 | 0.00 | 0.00 | 0.00 | 0.00 | 0.00 | 0.00 | 0.04 | 0.11 | 0.51 | 0.28 | 0.17 | 0.07 | 0.01 | 0.00 | 0.00 |
| Powdery_24h | 0.00 | 0.01 | 28.44 | 21.45 | 77.02 | 17.12 | 0.00 | 0.00 | 0.02 | 0.00 | 0.00 | 0.00 | 0.00 | 1.26 | 20.14 | 9.23 | 32.25 | 19.37 | 36.08 | 20.35 | 0.06 | 0.00 | 0.00 |
| Powdery_48h | 0.00 | 0.01 | 2.26 | 4.72 | 14.39 | 7.02 | 0.00 | 0.00 | 0.02 | 0.00 | 0.00 | 0.00 | 0.00 | 0.46 | 5.22 | 4.32 | 10.02 | 14.22 | 12.61 | 6.17 | 0.04 | 0.00 | 0.00 |
| Powdery_72h | 0.00 | 0.02 | 0.41 | 0.21 | 1.64 | 1.12 | 0.00 | 0.00 | 0.01 | 0.00 | 0.00 | 0.00 | 0.00 | 0.00 | 0.16 | 0.69 | 2.54 | 1.67 | 1.80 | 1.04 | 0.04 | 0.00 | 0.00 |
| Glenlea_Control_48pi | 0.00 | 0.00 | 0.00 | 0.00 | 0.00 | 0.10 | 0.00 | 0.00 | 0.00 | 0.00 | 0.00 | 0.00 | 0.00 | 0.00 | 0.00 | 0.03 | 0.10 | 0.11 | 0.06 | 0.23 | 0.05 | 0.00 | 0.00 |
| Glenlea_Fungus_48pi | 0.00 | 0.00 | 18.78 | 15.80 | 80.72 | 17.23 | 0.00 | 0.00 | 2.30 | 0.00 | 0.00 | 0.00 | 0.00 | 3.21 | 36.56 | 11.38 | 31.88 | 15.79 | 21.03 | 24.91 | 0.02 | 0.00 | 0.00 |
| Salamouni_Control_48pi | 0.00 | 0.00 | 0.00 | 0.00 | 0.00 | 0.18 | 0.00 | 0.00 | 0.00 | 0.00 | 0.00 | 0.00 | 0.00 | 0.00 | 0.00 | 0.15 | 0.50 | 0.35 | 0.35 | 0.46 | 0.05 | 0.00 | 0.00 |
| Salamouni_Fungus_48pi | 0.06 | 0.05 | 41.15 | 5.24 | ##### | 71.74 | 0.00 | 0.00 | 2.84 | 0.00 | 0.00 | 0.00 | 0.00 | 1.76 | 42.05 | 39.89 | 97.86 | 93.95 | ##### | 73.41 | 0.04 | 0.00 | 0.00 |
| non-innoculation | 0.00 | 0.00 | 0.00 | 0.03 | 0.09 | 0.10 | 0.00 | 0.00 | 0.00 | 0.00 | 0.00 | 0.00 | 0.00 | 0.00 | 0.04 | 0.11 | 0.51 | 0.28 | 0.17 | 0.07 | 0.01 | 0.00 | 0.00 |
| Stripe24h | 0.00 | 0.00 | 0.36 | 0.20 | 1.55 | 0.14 | 0.00 | 0.00 | 0.00 | 0.00 | 0.00 | 0.00 | 0.00 | 0.00 | 0.02 | 0.02 | 1.34 | 0.35 | 0.08 | 0.00 | 0.00 | 0.00 | 0.00 |
| Stripe48h | 0.00 | 0.00 | 0.00 | 0.00 | 0.12 | 0.07 | 0.00 | 0.00 | 0.00 | 0.00 | 0.00 | 0.00 | 0.00 | 0.01 | 0.04 | 0.06 | 0.63 | 0.19 | 0.38 | 0.05 | 0.01 | 0.00 | 0.00 |
| Stripe72h | 0.00 | 0.00 | 0.00 | 0.01 | 0.01 | 0.11 | 0.00 | 0.00 | 0.00 | 0.00 | 0.00 | 0.00 | 0.00 | 0.02 | 0.00 | 0.09 | 0.62 | 0.26 | 0.32 | 0.02 | 0.02 | 0.00 | 0.00 |
| Control_Root | 0.00 | 0.01 | 0.00 | 0.02 | 0.00 | 0.07 | 15.57 | 33.35 | 0.00 | 19.02 | 14.15 | 1.63 | 14.91 | 0.00 | 0.00 | 0.02 | 0.18 | 0.07 | 0.02 | 0.15 | 7.58 | 14.22 | 0.00 |
| Xt_Root | 0.01 | 0.12 | 0.05 | 0.00 | 0.03 | 0.38 | 15.50 | 28.56 | 0.00 | 19.20 | 14.36 | 1.47 | 14.43 | 0.00 | 0.06 | 0.12 | 0.25 | 0.05 | 0.11 | 0.03 | 6.05 | 14.99 | 0.00 |
| Control_Leaf | 0.00 | 0.00 | 0.00 | 0.00 | 0.00 | 0.00 | 0.00 | 0.00 | 0.00 | 0.00 | 0.00 | 0.00 | 0.00 | 0.00 | 0.00 | 0.00 | 0.12 | 0.00 | 0.01 | 0.02 | 0.00 | 0.00 | 0.00 |
| Xt_Leaf | 0.00 | 0.00 | 0.04 | 0.02 | 0.05 | 0.51 | 0.00 | 0.00 | 0.00 | 0.00 | 0.00 | 0.00 | 0.00 | 0.00 | 0.01 | 0.02 | 0.25 | 0.11 | 0.17 | 0.08 | 0.00 | 0.00 | 0.00 |
